# Tracing the invisible: Quantifying mirroring and embodied attunement in dyadic and triadic Dance Movement Therapy

**DOI:** 10.64898/2026.03.05.707373

**Authors:** S. Makris, B. Langley, R.M. Page, E. Perris, V. Karkou, V. Cazzato

**Affiliations:** Department of Theoretical and Applied Sciences, University eCampus, Novedrate (CO), Italy; Department of Sport and Physical Activity, Edge Hill University, Ormskirk, UK; Research Centre for Arts and Wellbeing, Faculty of Health, Social Care and Medicine, Edge Hill University, Ormskirk, UK; Department of Cognitive Sciences, Psychology, Education, and Cultural Studies, University of Messina, Messina (ME), Italy; Faculty of Health, Liverpool John Moores University, Liverpool, United Kingdom

**Keywords:** Dance Movement Therapy (DMT), Embodied Cognition, Therapist–Client Mirroring, Motion Capture Analysis, Intersubjective Synchrony

## Abstract

**Background:** Mirroring is a foundational method used in Dance Movement Therapy (DMT), assumed to foster empathy and therapeutic attunement, yet its embodied dynamics remain insufficiently studied. In this paper, we provide the first quantitative exploration of client-therapist mirroring across dyadic and triadic formats, examining how synchrony unfolded during a structured mirroring exercise in which participants alternated between leading and following roles.

**Methodology:** Using optical motion capture and time-series modelling, we quantified movement coordination in dyadic (female client–therapist; male client–therapist) and triadic (therapist with both clients) interactions.

**Results:** In dyadic tasks, the female client-therapist interaction was marked by tight temporal alignment, significant synchrony, robust predictive accuracy, and clear client-to-therapist influence, consistent with kinaesthetic empathy and affect-sensitive entrainment. By contrast, the male client–therapist dyad exhibited weaker and more delayed temporal coupling, alongside reduced phase synchronisation and fewer directional dependencies, despite comparable levels of interpersonal proximity. In the triadic task, temporal entrainment attenuated: therapist movement had few matching qualities to clients’ movement, yet recurrent synchrony with both clients persisted, suggesting a strategic shift from fine-grained entrainment to stable postural scaffolding under divided attention.

**Discussion:** These findings demonstrated that mirroring is not a uniform technique, but a family of embodied coordination modes flexibly recruited according to relational context and client expressivity. They align with theories of embodied simulation and affect attunement, implicating rapid motor resonance in dyadic entrainment and interoceptive–affective scaffolding in triadic stability. Clinically, the results underscore the need for training in flexible embodied strategies, split attention, and equitable allocation of attunement in group work. More broadly, they open a translational agenda linking kinematic synchrony to neural, interoceptive, and autonomic mechanisms, positioning mirroring as both an experiential hallmark and a measurable mechanism of change in embodied psychotherapy.

## Introduction

Dance Movement Therapy (DMT), also known as Dance Movement Psychotherapy (DMP) in the United Kingdom, is a form of psychotherapy in which movement and dance are used as primary media for communication and therapeutic change. It is defined as “a relational process in which client(s) and therapist use body movement and dance as an instrument of communication during the therapy process” (Association for Dance Movement Psychotherapy UK, 2025, p. 1). DMT is theoretically grounded in embodied cognition and emotion, which reject body–mind dualism and propose that affective, cognitive, and social processes are constituted through sensorimotor experience and action (Gallagher, 2005; Barsalou, 2008; Shapiro, 2011). Within this framework, movement is not merely expressive but plays a generative role in emotion regulation, interpersonal understanding, and relational presence. As such, DMT offers a naturalistic yet structured model system for investigating how embodied interaction contributes to psychological change.

A growing body of evidence supports the clinical relevance of DMT across a range of populations, including children, adults with depression, and older adults living with dementia (Bradt et al., 2015; Karkou et al., 2019, 2023; Lyons et al., 2018). Broader reviews of arts-based interventions further suggest beneficial effects of dance-based practices across mental health and neurological outcomes, although the limited number of DMT-specific trials prevents firm conclusions about treatment-specific mechanisms (de Witte et al., 2025). Despite this evidence base, the processes through which DMT exerts therapeutic effects remain poorly specified. Conceptual and qualitative work has identified candidate therapeutic factors—such as bodily awareness, movement expressivity, and non-verbal communication—but consistently highlights the therapeutic relationship as central, particularly kinaesthetic empathy supported by mirroring, movement dialogue, and interpersonal synchrony (de Witte et al., 2021; Koch, 2014). However, these constructs have rarely been operationalised in terms of observable, time-resolved movement dynamics.

Mirroring is a foundational method in DMT, typically involving the therapist’s responsive alignment with the client’s movement to foster kinaesthetic empathy, agency, and affective attunement. Neuroscientific accounts of action observation and embodied simulation provide a biologically plausible framework for this process, suggesting that observing and executing movement engage overlapping sensorimotor representations that support interpersonal understanding (Rizzolatti & Craighero, 2004; Gallese & Sinigaglia, 2011). Complementary research on interpersonal synchrony shows that temporal alignment of movement promotes affiliation, trust, and co-regulation, mediated by shared predictive and regulatory mechanisms (Hove & Risen, 2009; Mogan et al., 2017). However, empirical studies of mirroring in therapeutic contexts have relied predominantly on qualitative methods, limiting insight into its temporal structure, directionality, and flexibility. Moreover, existing research has focused almost exclusively on dyadic interactions, despite the frequent use of triadic and group formats in clinical practice, where therapists must distribute embodied attention across multiple partners under increased cognitive and relational load.

The present study addresses these gaps by applying optical motion capture and computational time-series analysis to therapist–client mirroring in both dyadic and triadic DMT interactions. Using a semi-structured mirroring task, we quantified the spatial and temporal organisation of movement coordination and examined how synchrony, directional influence, and recurrence patterns differed as a function of interaction format and client movement characteristics. By combining cross-correlation, phase synchronisation metrics, vector autoregression, and joint recurrence quantification analysis, we move beyond descriptive accounts to provide a quantitative characterisation of embodied attunement. In doing so, this work advances a mechanistic understanding of mirroring as a flexible family of coordination strategies rather than a uniform technique, with implications for psychotherapy research, social neuroscience, and the study of embodied interaction more broadly.

We hypothesised that dyadic mirroring would often exhibit stronger fine-grained temporal coordination than triadic mirroring, expressed as higher synchrony, shorter lagged coupling, and clearer directional dependencies between client and therapist movements. We further expected that, when fine-grained entrainment was present in dyads, directional influence would predominantly flow from client to therapist, consistent with affect-sensitive therapeutic following, whereas triadic interactions would show attenuated temporal entrainment due to divided attention and increased interactional complexity. We additionally hypothesised that this reduction in fine-grained synchrony in triadic mirroring would be accompanied by preserved or increased recurrence of shared movement states, reflecting a shift from rapid entrainment toward more stable, low-frequency postural or spatial scaffolding. Finally, we explored whether variation in client movement organisation would be associated with systematic differences in therapist coordination strategies across dyadic and triadic contexts, without assuming stable individual or gender-based effects but rather interaction-specific modulation of embodied attunement.

## Methods

### Design

The study used an idiographic, within-session comparative design to examine embodied coordination during therapist–client mirroring in Dance Movement Therapy (DMT). A single certified therapist interacted with two clients across two interaction formats—dyadic and triadic—allowing direct comparison of movement coordination as a function of interactional complexity while holding therapist, recording environment, and analytic framework constant. The design prioritised time-resolved analysis of movement dynamics within and across interaction formats rather than population-level inference, with dyadic and triadic conditions compared using complementary measures of temporal coordination, directional influence, and recurrence structure. The study was preregistered on the Open Science Framework (OSF); the registration is under embargo to preserve blind peer review.

### Participants

The study involved three participants: two clients (one female aged 34 and one male aged 31), both university students with little or no prior experience in DMT, and one female certified dance movement therapist. All participants were recruited voluntarily and provided written informed consent prior to taking part in the study. Participants were fully informed about the study’s purpose, procedures, and their right to withdraw at any stage without penalty and were assured of the confidentiality and anonymity of their data in accordance with ethical research guidelines. The therapist, who was unknown to the two client participants prior to the day of the data collection, had extensive professional experience in DMT and led the mirroring-based movement tasks. As the therapist’s movement data were included in the kinematic analyses, she was considered a participant for the purposes of data collection and analysis, while retaining a distinct functional role within the experimental design. Ethical approval for the study was obtained from the Edge Hill University institutional ethics board.

### Apparatus

Whole-body movement was recorded using a Qualisys optical motion capture system (Qualisys, Gothenburg, Sweden), equipped with ten infrared cameras (Oqus 3+) sampling at 200 Hz. The system tracked the three-dimensional positions of retroreflective markers with submillimetre accuracy, enabling high-resolution capture of full-body kinematics. Approximately 39 retroreflective markers were placed on key anatomical landmarks of all participants, including the head, trunk, pelvis, upper and lower limbs, feet, and hands, to allow modelling of major body segments in line with the marker configuration illustrated in Figure 1. To reduce model complexity and limit data dimensionality, a pragmatic modelling approach was adopted whereby whole-body movement was represented using the centre of mass (CoM) of the trunk segment, which served as a proxy for global body motion. This approach is consistent with prior movement synchrony research and is appropriate for capturing large-scale interpersonal coordination (Gaziv et al., 2017; Patoz et al., 2021; Tisserand et al., 2016). Movements of the trunk CoM were analysed along the three principal axes of motion: the medial–lateral (X; side-to-side) axis, the anterior–posterior (Y; forwards–backwards) axis, and the vertical (Z; up–down) axis. In addition to the motion capture system, two high-definition (HD) video cameras were used to provide supplementary qualitative recordings of the sessions. One camera was positioned from the therapist’s perspective to capture clients’ movements and responses, while the second camera was placed from the client’s perspective to record the therapist’s mirroring actions. This dual-camera configuration provided comprehensive visual documentation of the dyadic and triadic interactions, complementing the quantitative kinematic data. Details of motion capture data processing and analytical procedures are reported in a dedicated section prior to the statistical analyses.

**Figure 1.**
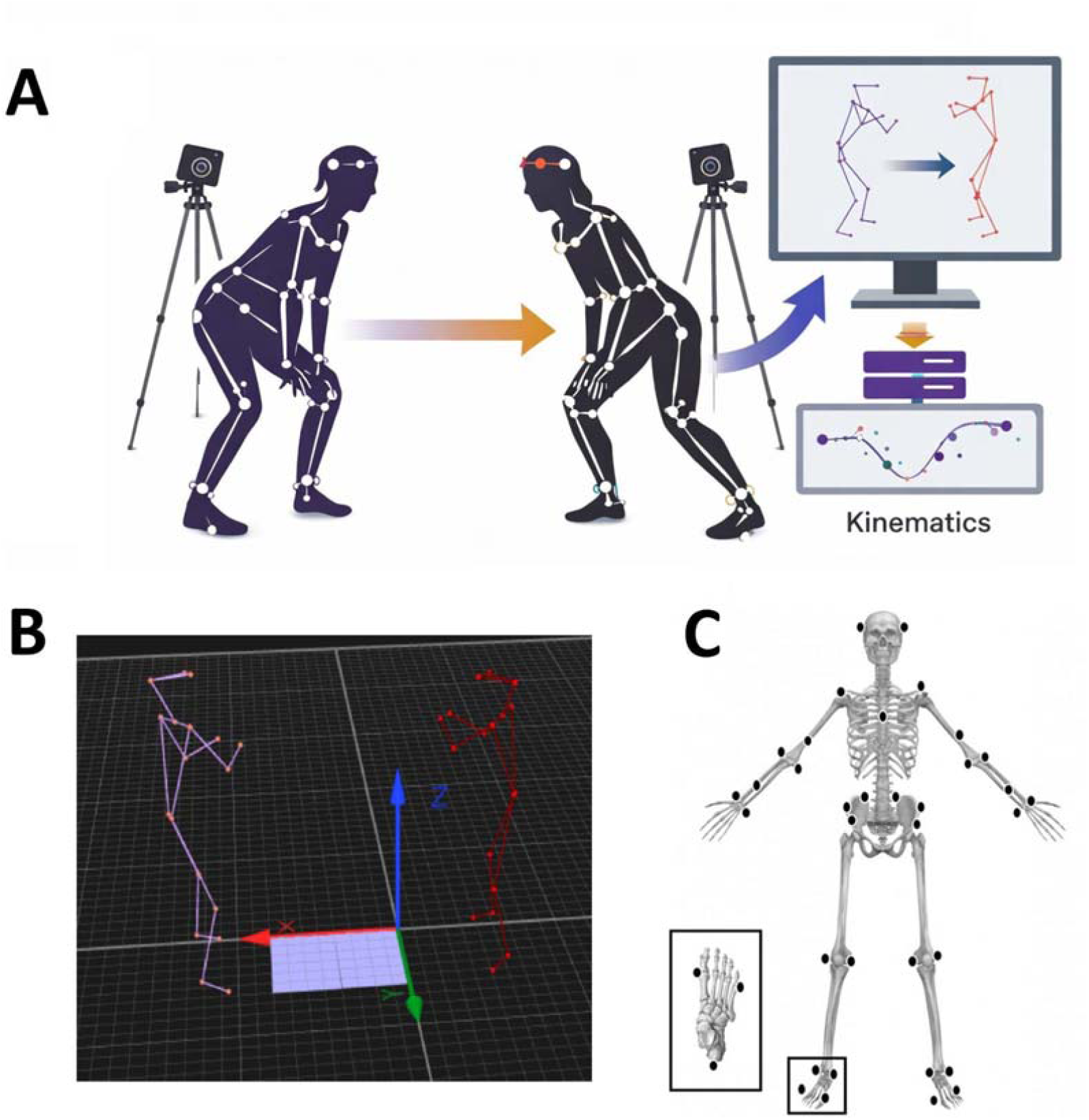
Example of the experimental setup during a dyadic dance movement session with marker-based motion capture. A) Schematic representation of the dyadic dance interaction and motion-capture data processing pipeline. B) Three-dimensional skeletal reconstruction of the two participants in the shared coordinate space, used for kinematic analyses. C) Marker placement for motion capture. Markers located on the central, posterior aspect of the cranium and at 7th cervical vertebrae not shown on the image.

### DMT tasks

The study involved three structured DMT tasks conducted in a controlled environment, with each client engaging in a series of interactions with the therapist under two conditions: dyadic (one-to-one interaction) and triadic (interaction with both clients simultaneously). Each dyadic task lasted approximately 15 minutes, while the triadic task lasted around 20 minutes. The therapist led the tasks with a strong emphasis on mirroring exercises, aiming to foster emotional attunement and enhance body awareness through embodied empathic communication. She followed a structured approach, incorporating both spontaneous and guided movement sequences. Clients were encouraged to engage in natural and expressive movement while the therapist mirrored their actions with the goal of deepening their awareness of internal states and external expressions. In the dyadic tasks, the therapist closely observed each client’s non-verbal communication, responding to their movements with attuned gestures that sought to demonstrate understanding and provide a reflective space for self-exploration.

With the male client, the therapist engaged in a series of gradual and rhythmic movements, including mirroring finger movements, tapping, and body scanning exercises. The therapist’s responses were aimed at externalizing the client’s internal state and providing a mirror to support self-awareness. The therapist often invited the male client to explore sensations through structured movement prompts, such as tapping across the body and shifting between open and closed body positions, to facilitate an embodied understanding of emotional states.

The female client’s interactions involved more variation in effort qualities and smoother transitions between movement patterns, which required the therapist to make frequent and adaptive adjustments. The therapist mirrored swaying, rocking, and self-touch movements, encouraging the client to engage with areas of bodily sensation connected to emotion. Throughout the task, the therapist employed techniques such as guided breath awareness, tactile stimulation through stroking, and the use of metaphor to facilitate exploration of emotional states.

During the triadic task, the therapist faced the additional complexity of synchronising with both clients simultaneously. The task began with an invitation for both clients to share an initial movement reflecting their current state. The therapist then introduced rhythmic elements, guiding the group through collective and individual movement explorations. Clients were encouraged to follow shared rhythmic patterns and experiment with variations in speed and tempo. This exercise was designed to promote group cohesion while allowing individual expressions to emerge within the shared movement framework.

Throughout the tasks, intentional pauses were incorporated to facilitate 3D motion capture recordings, with video recordings capturing qualitative insights and supporting the quantitative analysis of movement data. The therapist alternated between active mirroring and neutral observation stances to assess clients’ ability to engage with their movements independently.

The therapist’s interventions were guided by principles of embodied empathic communication, with a focus on demonstrating attunement to clients’ movements, facilitating new ways of engaging with bodily sensations, and encouraging exploration of unconscious experiences through movement. The recorded activities provided valuable insights into the dynamic interplay between the therapist’s mirroring responses and clients’ evolving movement patterns.

### Data Processing

Mirroring phases were therapist-initiated as an explicit component of the intervention protocol and were not inferred from kinematic features. The onset of mirroring was indicated in situ by the therapist using a pre-agreed unobtrusive visual cue (“secret sign”) visible in the video recordings. Following data collection, the therapist reviewed the session videos to confirm the onset and offset of the mirroring phase and to verify correspondence with the intended therapeutic procedure. Independent validation was provided by an external UK-based qualified Dance Movement Therapy (DMT) therapist, not otherwise involved in the study and blinded to the study hypotheses, who reviewed the relevant video excerpts and confirmed the presence of therapist-led mirroring within the identified intervals. Kinematic analyses were restricted exclusively to these therapist-indicated and independently validated mirroring segments. No segments were selected, modified, or excluded on the basis of kinematic properties, synchrony metrics, or post hoc inspection of movement data.

Data analysis focused on short segments of movement-based interaction extracted from full Dance Movement Therapy sessions under both dyadic and triadic conditions. These segments corresponded to discrete periods during which the therapist explicitly engaged in mirroring the clients’ movements. Each analysed segment had a duration of approximately 40–45 seconds, providing a standardised temporal window for assessing interpersonal coordination while maintaining sustained engagement by all participants. For each participant and condition, three-dimensional centre-of-mass (CoM) trajectories of the trunk segment were extracted and analysed along the medio–lateral (ML), antero–posterior (AP), and vertical (V) axes. Spatial organisation of the interaction was further characterised by computing the Euclidean distance between therapist and client CoM trajectories. Where required by specific analytical procedures, CoM time series were normalised within participants and trials to facilitate comparability while preserving the temporal structure of the movement signals. Because mirroring phases were embedded as a discrete element of the therapeutic protocol, a single continuous mirroring episode was available and analysed for each condition, yielding an idiographic, case-comparative assessment rather than population-level inference. Full details of motion-capture reconstruction, filtering, coordinate transformations, and preprocessing procedures are reported in the Supplementary Materials.

### Data Analysis

Temporal coordination between therapist and client was first assessed using cross-correlation analysis of centre-of-mass (CoM) trajectories. Cross-correlation functions were computed within a ±5 s lag window, and the peak correlation coefficient (r) was used to identify the temporal offset at which movements were most strongly aligned. Positive lag values indicated therapist-following (client-leading) dynamics, whereas negative lag values indicated therapist-leading dynamics. Correlation magnitudes are reported descriptively to aid interpretation.

Phase-based movement synchrony was quantified using a Synchronisation Index (SI), ranging from 0 (no phase alignment) to 1 (perfect phase-locking). Statistical significance of SI values was assessed using circular-shift permutation testing, with observed values considered significant if they exceeded the 95th percentile of the corresponding null distribution.

Spatial organisation of the interaction was assessed using inter-centre-of-mass distance. Three-dimensional Euclidean distances between therapist and client CoM trajectories were computed over time and expressed relative to the initial inter-CoM separation to facilitate comparison across conditions. Descriptive statistics were used to characterise the magnitude and temporal stability of interpersonal spacing.

Directional temporal dependencies were examined using Granger causality analyses conducted on down-sampled, z-standardised CoM trajectories. Lag order was selected using the Akaike Information Criterion, and Granger tests evaluated whether past values of one partner’s movement significantly improved prediction of the other partner’s movement beyond autoregressive effects alone. Results are reported as F-statistics with associated p-values.

Where appropriate, correction for multiple comparisons was applied within method-specific families of tests. Full details of predictive modelling, multivariate analyses, preprocessing steps, and statistical controls are provided in the Supplementary Materials.

## Results

### Dyadic mirroring with female client

Cross-correlation analyses revealed strong temporal coupling between the female client and therapist, with predominantly client-leading dynamics (see also Figure 2). In the antero–posterior axis, peak coupling was strong (r = .749) and occurred at a short negative lag (−0.73 s), indicating that the therapist’s movements closely followed the client’s forward–backward dynamics. Similarly, vertical movement showed very strong coupling (r = .977 at −0.39 s), reflecting highly consistent mirroring within a sub-second temporal window. In the medial–lateral axis, coupling was characterised by a pronounced anti-phase relationship. The strongest association was negative (r = −.706 at −0.34 s), indicating that therapist lateral movements were temporally aligned with, but opposite in direction to, those of the client (e.g., leftward displacement of the client accompanied by rightward displacement of the therapist). Zero-lag correlations further confirmed concurrent coupling in the antero–posterior (r = .715, p = .007) and vertical (r = .967, p = .001) dimensions, whereas the medial–lateral axis showed a significant negative concurrent association (r = −.628, p = .017).

**Figure 2.**
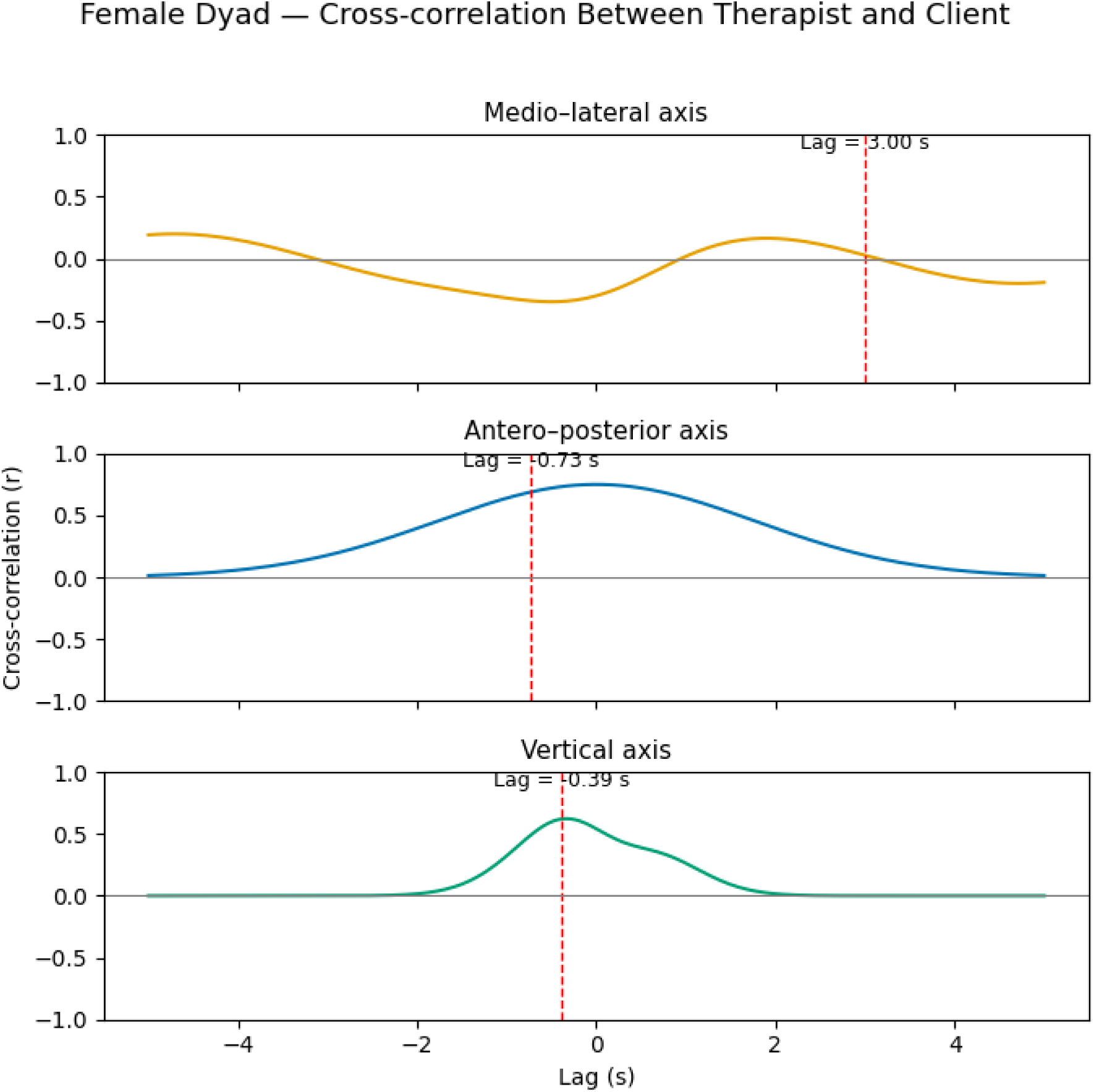
Cross-correlation functions between therapist and female client centre-of-mass trajectories along the medio–lateral (ML), antero-posterior (AP), and vertical (V) axes in the dyadic condition. Dashed vertical lines indicate the lag at which the maximum absolute cross-correlation coefficient was observed within a ±5 s window. Positive lags indicate therapist-following dynamics, whereas negative lags indicate therapist-leading dynamics.

The time-series analysis revealed that medio–lateral synchronisation was low (SI = 0.338) and did not exceed the permutation-derived significance threshold (p = .475). Similarly, antero–posterior synchronisation was low (SI = 0.292) and non-significant (p =.305). In contrast, vertical synchronisation was significantly greater than chance (SI = 0.626, p = .032), exceeding the 95th percentile of the permutation null distribution. Statistically significant phase-locking between therapist and client was therefore observed only along the vertical axis across the 0.1–3 Hz frequency range (see also Figure 3).

**Figure 3.**
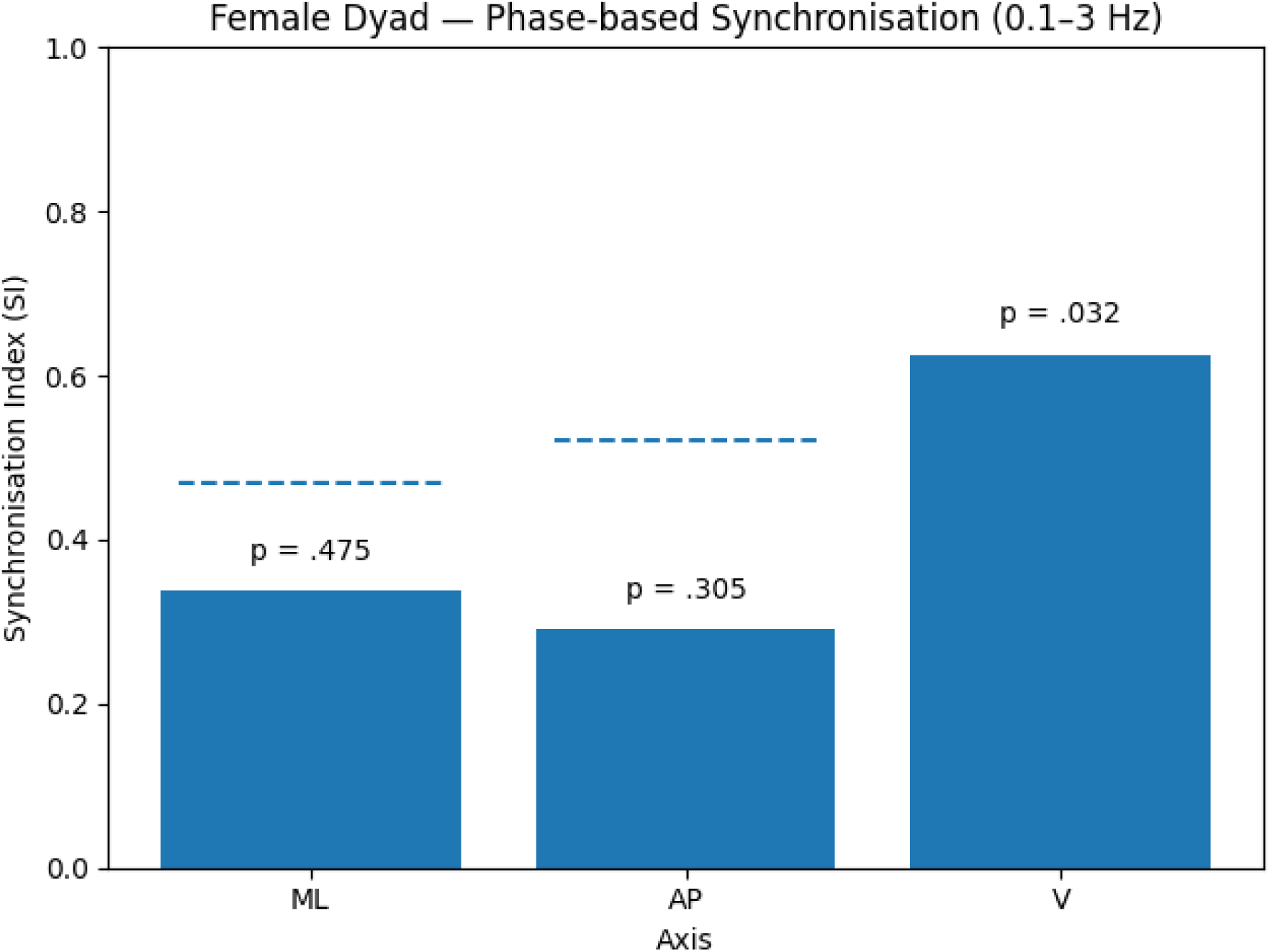
Phase-based synchronisation between therapist and female client in the dyadic condition. Synchronisation Index (SI) values were computed from z-standardised centre-of-mass trajectories band-pass filtered between 0.1 and 3 Hz. Dashed lines indicate permutation-derived significance thresholds; p-values are shown above bars.

Having characterised temporal coordination between therapist and client, spatial organisation of the interaction was examined using normalised inter-centre-of-mass (CoM) distance. When expressed relative to the initial separation, the mean normalised inter-CoM distance was 1.042 (SD = 0.416), with values ranging from 0.238 to 2.306. These values indicate that, on average, therapist–client distance remained close to the initial separation, while exhibiting substantial variability over time.

Predictive coupling analyses using nonlinear regression yielded convergent evidence for directional asymmetry in dyadic interaction, with therapist movement more predictable from client dynamics in the horizontal planes; full Random Forest results are reported in the Supplementary Materials.

Granger causality analyses on z-standardised centre-of-mass (CoM) trajectories revealed axis-specific patterns of directional influence between client and therapist (see also Figure 4). Along the medio–lateral (ML) axis, significant bidirectional effects were observed, with the client’s past dynamics predicting therapist movement, F(7, 988) = 6.62, p < .001, and a weaker but significant reverse influence, F(7, 988) = 2.14, p = .038. In the antero–posterior (AP) axis, a clear directional asymmetry emerged: client movement significantly predicted therapist movement, F(6, 991) = 3.34, p = .003, whereas the therapist-to-client pathway was not significant, F(8, 985) = 0.74, p = .655. In the vertical axis, strong bidirectional coupling was evident, with significant effects in both directions (client → therapist: F(7, 988) = 13.95, p < .001; therapist → client: F(6, 991) = 3.09, p = .005). All reported effects survived FDR correction within condition. Overall, Granger causality analyses indicated a predominant client-to-therapist influence across movement dimensions, alongside reciprocal coupling in the ML and vertical axes.

**Figure 4.**
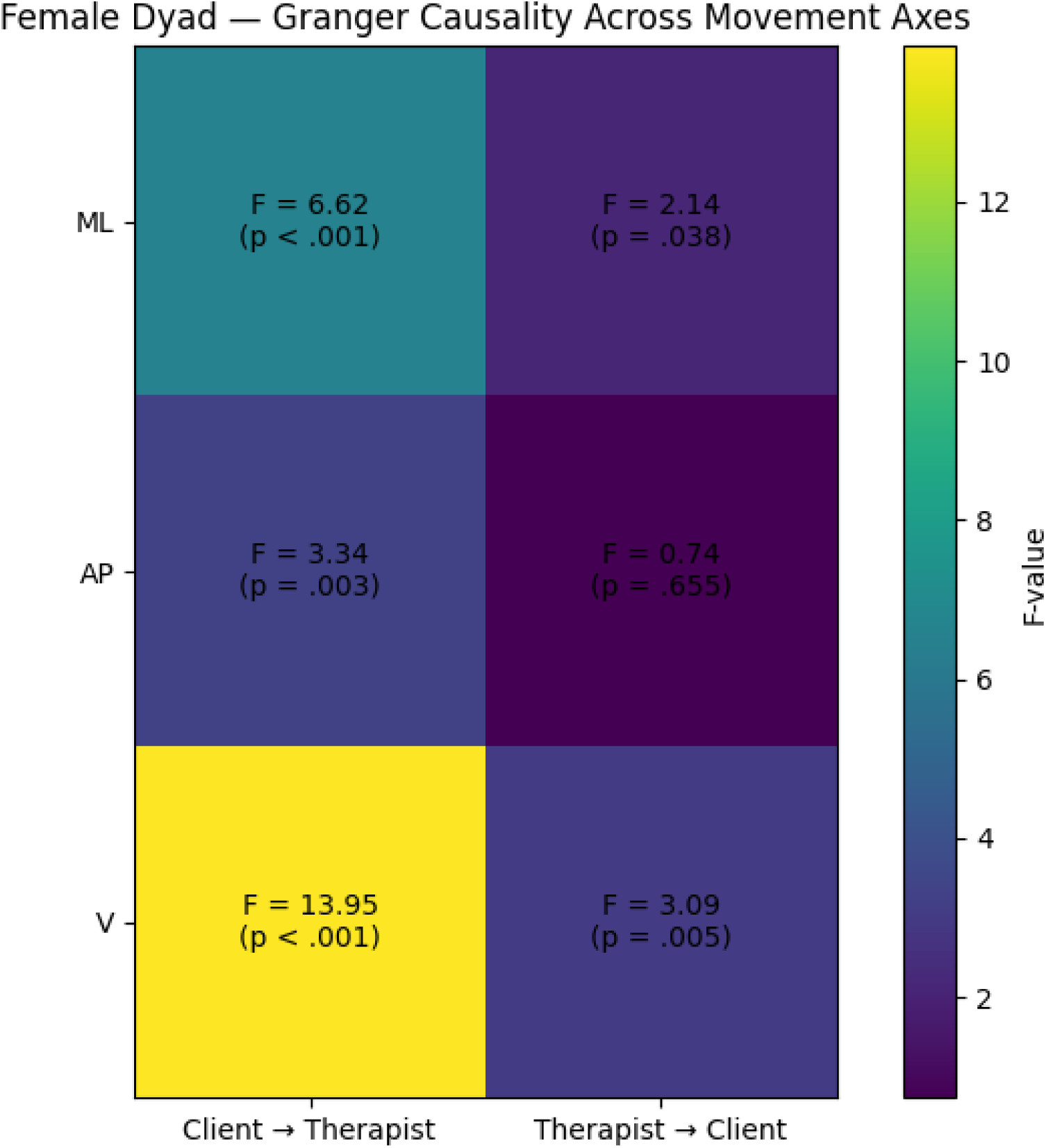
Heatmap showing Granger causality results for the female dyad across movement axes. Cells display F-statistics from Granger causality tests assessing directional influence between client and therapist centre-of-mass trajectories, with corresponding p-values indicated in parentheses. Colour intensity reflects the magnitude of the F-statistic. Significant effects (after FDR correction within condition) indicate that past values of one partner’s movement significantly improved prediction of the other partner’s subsequent movement beyond autoregressive effects alone. ML = medio–lateral; AP = antero–posterior; V = vertical.

### Dyadic mirroring with male client

The same analytical approach was applied to the dyadic mirroring interaction involving the male client. Along the medio–lateral (ML) axis, the cross-correlation function peaked at a low magnitude (r = .217) with an optimal lag of 1.89 s, indicating weak client-leading coordination. In contrast, the antero–posterior (AP) axis exhibited a stronger peak correlation (r = .515) at −2.10 s, indicating therapist-leading dynamics. Similarly, vertical (V) axis coupling peaked at r = .511 with a lag of −1.97 s, again reflecting therapist-leading temporal coordination. Negative correlation peaks were also observed in the ML (r = −.172) and V (r = −.249) axes, indicating intermittent anti-phase relationships between therapist and client movements. Overall, temporal coupling in the male dyad was weak and delayed in the ML axis, but stronger in the AP and vertical axes, where coordination was predominantly therapist-led.

Phase synchronisation analysis revealed low and non-significant synchrony across all axes in the male dyad. In the medio–lateral (ML) axis, synchronisation was low (SI = 0.076) and did not exceed the permutation-derived significance threshold (p = .891). Similarly, synchronisation in the antero–posterior (AP) axis was low (SI = 0.145, p = .741). The vertical axis showed comparatively higher SI values (SI = 0.257), but this effect also did not reach statistical significance (p = .541). Thus, no statistically significant phase-locking was observed in any movement dimension for the male dyadic interaction.

Spatial organisation of the male dyad was assessed using normalised inter-centre-of-mass (CoM) distance. When expressed relative to the initial separation, the mean normalised inter-CoM distance was 0.762 (SD = 0.362), with values ranging from 0.149 to 2.355. These values indicate that, relative to the initial configuration, therapist–client distance frequently decreased during the interaction, while also exhibiting substantial variability over time.

Nonlinear predictive modelling further indicated a reversal of directional asymmetry in the male dyad relative to the female dyad, with therapist movements more predictive of client dynamics in the horizontal planes; full Random Forest results are reported in the Supplementary Materials.

Granger causality analyses for the male dyad revealed no evidence of directional influence in the horizontal planes. Along the medio–lateral (ML) axis, neither direction showed significant Granger causality (client → therapist: F(6, 991) = 0.41, p = .869; therapist → client: F(8, 985) = 0.63, p = .755). Similarly, no significant effects were observed in the antero–posterior (AP) axis (client → therapist: F(8, 985) = 1.45, p = .173; therapist → client: F(8, 985) = 1.05, p = .399). By contrast, in the vertical (V) axis a unidirectional influence from client to therapist was observed, with the client’s past dynamics significantly improving prediction of the therapist’s subsequent movements (F(6, 991) = 3.22, p = .0039). The reverse pathway did not reach significance (F(7, 988) = 0.40, p = .900). All reported effects survived FDR correction within condition. Overall, Granger causality analyses indicated selective client-to-therapist influence confined to the vertical movement dimension.

### Comparative Results: Female vs Male Dyad

Direct comparison of client–therapist mirroring across the female and male dyadic interactions revealed clear differences in temporal coordination, phase synchronisation, and directional coupling (see also Table 1). In the female dyad, therapist movements showed close temporal alignment with those of the client, as evidenced by short-lag cross-correlations, significant vertical phase synchronisation, and consistent client-to-therapist directional influences across multiple movement dimensions. Granger causality analyses indicated predominant client-led directionality, with reciprocal coupling confined to specific axes.

**Table 1.** Summary of key statistical indicators of therapist–client mirroring in the female and male dyadic conditions. Reported metrics include peak lagged cross-correlation values (within a ±5 s window), phase synchronisation indices (SI) with permutation-derived significance levels, mean absolute inter-centre-of-mass (CoM) distance expressed in metres (with coefficient of variation, CV), Random Forest predictive performance (out-of-sample R² from three-block time-ordered cross-validation, reported separately for each prediction direction), and Granger causality results indicating significant directional temporal dependencies. Negative lag values indicate therapist-leading dynamics, whereas positive values indicate client-leading dynamics. Absolute distance values reported here are complemented by normalised distance measures relative to initial separation in the Supplementary Materials, which provide full time-resolved spatial profiles.

| Metric | Female Dyad | Male Dyad | Summary |
| --- | --- | --- | --- |
| *Cross-correlation*<br>(*peak lag*) | AP: -0.73 s; V: -0.39 s;<br>ML: +3.00 s | AP: -2.10 s; V: -1.97 s;<br>ML: +1.89 s | Shorter, more<br>structured lags in<br>female dyad |
| *Synchronisation Index*<br>(*SI*) | V axis significant (SI =<br>0.626, $p = .032$) | All axes non-significant<br>(SI $\leq 0.257$) | Phase-locking<br>only in female dyad |
| *Mean inter-CoM*<br>*distance* | 0.454 m (CV = 0.399) | 0.440 m (CV = 0.475) | Comparable<br>proximity; higher<br>variability in male<br>dyad |
| *Granger causality* | Client→therapist in ML,<br>AP, V; reciprocal in ML, V | Client→therapist in V only | Directionality<br>broader in female<br>dyad |

In contrast, the male dyad exhibited weaker and more delayed temporal organisation. Cross-correlation peaks were generally longer-lagged and more frequently therapist-led, phase synchronisation indices did not reach statistical significance in any axis, and Granger causality effects were restricted to a unidirectional client-to-therapist influence in the vertical dimension only. Spatial organisation, assessed using normalised inter-centre-of-mass distance, was broadly comparable across dyads in terms of mean proximity. However, the male dyad exhibited greater variability in interpersonal distance over time, whereas the female dyad showed more stable spatial organisation. Together, these findings indicate that client-led temporal structure was more consistently expressed—and more systematically mirrored—during the female dyadic interaction than in the male dyad. These differences are interpreted as interaction-specific coordination patterns arising within a brief, task-constrained context, rather than as stable individual characteristics or indices of therapeutic efficacy.

### Triadic mirroring

Although analyses in this section are reported pairwise for clarity and comparability with dyadic conditions, the triadic task constitutes a coupled three-person system in which movement dynamics are mutually interdependent. Pairwise metrics should therefore be interpreted as local projections of a higher-dimensional group process rather than as independent therapist–client exchanges.

### Female client

In the triadic condition involving the female client, cross-correlation patterns differed across movement dimensions. Along the medio–lateral (ML) axis, coupling was weak and temporally delayed, with low-magnitude correlations occurring at longer lags. Similarly, coordination in the antero–posterior (AP) axis was limited, with delayed and predominantly negative associations indicating weak alignment. By contrast, the vertical (V) axis exhibited strong coupling at near-zero lag. Peak correlation occurred at −0.09 s (r = .747), indicating that the client’s vertical movements consistently preceded those of the therapist by a fraction of a second. Overall, cross-correlation analyses indicated that, in the triadic setting, client–therapist temporal alignment was concentrated in the vertical dimension, whereas coordination in the horizontal planes was weak and delayed. Full axis-specific cross-correlation values and lag distributions are reported in the Supplementary Materials.

Phase synchronisation patterns in the triadic condition involving the female client revealed no statistically significant phase-locking across movement dimensions. In the medio–lateral (ML) axis, synchronisation was low (SI = 0.242, p = .687), while the antero–posterior (AP) axis showed minimal phase alignment (SI = 0.035, p = .969). The vertical (V) axis exhibited comparatively higher SI values (SI = 0.308), but this effect did not exceed the permutation-derived significance threshold (p = .241). Thus, no statistically significant phase-based coupling between therapist and client was observed in the triadic condition across any axis.

Spatial organisation of the therapist–female client interaction in the triadic condition was assessed using normalised inter-centre-of-mass (CoM) distance. When expressed relative to the initial separation, the mean normalised inter-CoM distance was 0.809 (SD = 0.278), with values ranging from 0.110 to 1.528. These values indicate that, relative to the initial configuration, therapist–client distance was frequently reduced during the interaction, while also exhibiting variability over time.

Nonlinear predictive modelling in the triadic condition yielded axis-specific patterns of directional predictability that were broadly consistent with the inferential analyses; full Random Forest results are reported in the Supplementary Materials.

Granger causality analyses on z-standardised centre-of-mass (CoM) trajectories in the triadic condition revealed axis-specific patterns of directional influence between the female client and therapist. Along the medio–lateral (ML) axis, neither direction reached statistical significance (female → therapist: F(7, 988) = 1.64, p = .119; therapist → female: F(8, 985) = 1.02, p = .420). Similarly, no evidence of directional influence was observed in the antero–posterior (AP) axis (female → therapist: F(7, 988) = 0.82, p = .568; therapist → female: F(7, 988) = 0.90, p = .509). By contrast, the vertical (V) axis exhibited a significant unidirectional effect from the female client to the therapist (F(7, 988) = 3.48, p = .0011), whereas the reciprocal pathway did not reach significance (F(8, 985) = 1.51, p = .150). All reported effects survived FDR correction within condition. Overall, Granger causality analyses indicated selective client-to-therapist directional influence confined to the vertical movement dimension in the triadic interaction.

### Male client

In the triadic condition involving the male client, cross-correlation patterns varied across movement dimensions. Along the medio–lateral (ML) axis, coupling was weak and temporally delayed, with low-magnitude correlations occurring at longer lags. Similarly, coordination in the antero–posterior (AP) axis was limited, with predominantly negative and near-zero associations indicating weak or inconsistent temporal alignment. By contrast, the vertical (V) axis exhibited strong coupling at a short positive lag. Peak correlation occurred at +0.43 s (r = .754), indicating that therapist vertical movement slightly followed that of the male client on a sub-second timescale. Overall, cross-correlation analyses indicated that, in the triadic interaction with the male client, client–therapist temporal alignment was concentrated in the vertical dimension, whereas coordination in the horizontal planes was weak and delayed. Full axis-specific cross-correlation values and lag distributions are reported in the Supplementary Materials.

Phase synchronisation analyses in the triadic condition involving the male client revealed no statistically significant phase-locking across movement dimensions. In the medio–lateral (ML) axis, synchronisation was very low (SI = 0.050, p = .938). The antero–posterior (AP) axis exhibited higher SI values (SI = 0.217), but this effect did not exceed the permutation-derived significance threshold (p = .453). The vertical (V) axis showed the largest SI values (SI = 0.305), yet synchronisation remained non-significant (p = .508). Thus, no statistically significant phase-based coupling between therapist and client was observed in the triadic interaction across any axis.

Spatial organisation of the therapist–male client interaction in the triadic condition was assessed using normalised inter-centre-of-mass (CoM) distance. When expressed relative to the initial separation, the mean normalised inter-CoM distance was 0.753 (SD = 0.309), with values ranging from 0.053 to 1.409. These values indicate that, relative to the initial configuration, therapist–client distance was frequently reduced during the interaction, while also exhibiting variability over time.

Nonlinear predictive modelling in the triadic condition involving the male client yielded axis-specific patterns of directional predictability that were broadly consistent with the inferential analyses; full Random Forest results are reported in the Supplementary Materials.

Granger causality analyses on z-standardised centre-of-mass (CoM) trajectories for the male client in the triadic condition revealed axis-specific patterns of directional influence. Along the medio–lateral (ML) axis, no significant Granger effects were observed in either direction (male → therapist: F(6, 991) = 0.96, p = .449; therapist → male: F(7, 988) = 0.55, p = .796). Similarly, the antero–posterior (AP) axis showed no evidence of directional influence (male → therapist: F(7, 988) = 1.49, p = .168; therapist → male: F(8, 985) = 1.25, p = .267). By contrast, the vertical (V) axis exhibited significant bidirectional Granger effects, with the male client’s past dynamics predicting the therapist’s subsequent movements (F(7, 988) = 2.42, p = .019) and the therapist’s dynamics also significantly predicting the client’s movements (F(7, 988) = 2.34, p = .023). All reported effects survived FDR correction within condition. Overall, Granger causality analyses indicated reciprocal directional influence confined to the vertical movement dimension in the triadic interaction.

### Multivariate Modelling of Therapist Mirroring in Triadic Task

To assess therapist coordination with both clients simultaneously in the triadic task, multivariate analyses were conducted to complement the pairwise results. Across movement dimensions, therapist movement was primarily characterised by strong autoregressive structure, with no reliable evidence that either client’s prior dynamics systematically predicted the therapist’s subsequent movements when modelled jointly. Consistent with this pattern, analyses of shared movement structure indicated the presence of temporally organised recurrence between therapist and clients in the vertical dimension, but without clear differentiation between therapist–female and therapist–male dyads. Full multivariate modelling results, recurrence metrics, and associated figures are reported in the Supplementary Materials.

## Discussion

The present findings offer new empirical evidence for the embodied mechanisms of client-therapist mirroring in DMT by directly comparing dyadic and triadic tasks. Through optical motion capture and computational modelling, we demonstrate that client-therapist interactions are shaped not by a singular act of mimicry but by a spectrum of coordination modes that are flexibly recruited in response to relational context and expressive salience. In dyads, and particularly in the female client-therapist interaction, coordination manifested as tight temporal coupling and strong directional predictability, while in the triadic task temporal entrainment weakened yet recurrent postural synchrony remained pronounced. This differentiation between entrainment and structural stability speaks to the multicomponent nature of embodied alignment and provides a quantitative framework for describing how therapists adapt their mirroring behaviour across contexts (Koole & Tschacher, 2016; Samaritter & Payne, 2013; McGarry & Russo, 2011).

The dyadic findings, particularly those involving the female client, converge with a broader literature demonstrating that temporal alignment in movement is associated with interpersonal attunement, empathy, and rapport (Hove & Risen, 2009; Valdesolo & DeSteno, 2011), and with psychotherapeutic research indicating that bodily coordination predicts therapeutic alliance and treatment outcomes (Ramseyer & Tschacher, 2011; Tschacher & Meier, 2020). In the present data, this alignment was expressed quantitatively through significant phase synchronisation, strong predictive coupling whereby therapist behaviour was reliably forecast from client kinematics, and directional temporal asymmetries indicating client-leading dynamics in Granger analyses. Together, these results provide convergent evidence that the therapist’s movement responses were systematically contingent on the client’s ongoing motor behaviour, rather than reflecting generic or self-driven synchrony. Within DMT, such patterns are often conceptualised in terms of kinaesthetic empathy (Koch & Fischman, 2011; Tortora, 2009), whereby the therapist’s embodied responses are shaped by the client’s expressive signals. More broadly, however, the present findings can also be situated within the person-centred concept of empathy (Rogers, 1951) and developmental and interactional frameworks such as Stern’s construct of affect attunement (Stern, 1985, 2010). The latter emphasises matching of temporal structure, intensity, and dynamic contour as a mechanism for communicating recognition of another’s internal state. From this perspective, the observed client-led temporal organisation provides a mechanistic account of how embodied responsiveness may emerge in therapeutic interaction, independently of any single theoretical tradition.

By contrast, the therapist–male client dyad exhibited weaker and more temporally delayed coupling, characterised by non-significant phase synchronisation and reduced directional predictability. A plausible interpretation of this asymmetry, consistent with established principles in DMT and nonverbal communication research, is that the temporal clarity and rhythmic organisation of client movement afford differential opportunities for embodied mapping. When movement is rhythmically structured and affectively coherent, it provides a legible temporal signal that can be more readily mirrored within the therapist’s motor system, facilitating simulation-based resonance and temporal alignment. Conversely, when movement is less clearly structured or dynamically ambiguous, entrainment may be more fragile and less amenable to fine-grained temporal coordination (Christov-Moore et al., 2014; Fischer & LaFrance, 2015). Clinical accounts within DMT have long emphasised that, under such conditions, mirroring is typically modulated rather than abandoned, with therapists shifting from micro-timed following towards strategies that prioritise simplified rhythmic contours, postural correspondence, or spatial alignment (Dosamantes-Beaudry, 2003; Samaritter & Payne, 2013). The present findings are consistent with this pattern: despite the absence of statistically significant phase-based synchrony, the maintenance of stable spatial proximity suggests that the therapist regulated the interaction through spatial scaffolding, supporting the relational frame without relying on high-cost temporal entrainment. Population-level differences in nonverbal expressivity and decoding have been documented (Hall, 2006; Fischer & LaFrance, 2015) and may have contributed to the observed asymmetries without determining individual interactional dynamics. In this context, the female client’s more rhythmically and affectively legible movement dynamics were associated with rapid embodied tracking, whereas the male client’s less temporally structured movement profile coincided with greater reliance on spatial forms of therapeutic containment.

The triadic task provides a more complex picture that extends beyond dyadic resonance. Cross-correlation analyses again highlighted low-frequency movement as a principal channel of coupling, with particularly consistent effects in the vertical dimension, in line with evidence that slow postural dynamics support interpersonal coordination (Zivotofsky & Hausdorff, 2007). At the same time, side-to-side organisation—a common stabilising strategy in DMT group work—also appeared to contribute to maintaining shared structure, even when precise temporal alignment was attenuated. Unlike in the dyadic condition, however, phase-locking did not reach statistical significance in any axis, suggesting that triadic coordination relied less on fine-grained temporal synchrony and more on broader structural organisation. Pairwise Granger causality analyses nonetheless indicated directed temporal dependencies—female client to therapist, and bidirectional exchange with the male client—but these effects were substantially attenuated in the multivariate VAR once covariance across the full three-person system was modelled. Such attenuation is well documented in time-series analyses when apparent bivariate dependencies share variance with a third partner or with strong self-dependence, underscoring the importance of modelling the interaction as a coupled system rather than as independent dyads. Importantly, joint recurrence quantification analysis (jRQA) revealed high determinism and laminarity across both therapist–client pairings, slightly higher in the therapist–male dyad. This indicates that, even in the absence of precise phase-locking, clients repeatedly returned to and sustained shared kinematic configurations over time. Clinically, this pattern aligns with descriptions of divided-attention work in DMT, in which therapists stabilise the group field through simple, durable movement anchors such as rocking from side to side —involving side-to-side weight shifts and gentle vertical rise-and-fall—so that inclusion and co-regulation are preserved across partners (Payne, 2017). The marginally higher recurrence observed with the male client may reflect a strategic emphasis on maintaining structural support for the less temporally expressive partner, thereby sustaining equitable embodied engagement within the triadic configuration (Meekums, 2020), even as the female client continued to exert greater moment-to-moment directional influence.

These behavioural dissociations dovetail with recent work in dance, music and social neuroscience. Novembre and colleagues show that musical/movement synchrony recruits partially distinct neural signatures: high-frequency phase entrainment engages sensorimotor coupling in premotor and parietal cortices, whereas slower, recurrent joint states involve shared interoceptive–affective regulation via anterior insula and cingulate (Novembre & Keller, 2014; Novembre et al., 2017). In line with this interpretation, the dyadic female interaction pattern may be plausibly characterised by an AON–dominant form of affect-sensitive entrainment, consistent with accounts emphasising sensorimotor resonance, action mirroring, and shared affective tuning during interpersonal coordination (Rizzolatti & Craighero, 2004; Keysers et al., 2010). Within dyadic contexts, such entrainment is likely supported by a body-centred mode of engagement, in which participants dynamically integrate exteroceptive and interoceptive signals to sustain mutual coordination. By contrast, the triadic configuration appears to shift attentional and computational resources towards exteroceptive, visually mediated processing, reflecting the need to monitor multiple agents and manage increased social exposure. Under these conditions, interoceptive focus may be attenuated or strategically downregulated, favouring outward-oriented coordination and behavioural regulation rather than enhanced interoceptive scaffolding or affect regulation per se. This interpretation is consonant with embodied simulation theory (Gallese, 2005; Gallese & Sinigaglia, 2011), which posits that perceiving another’s movement recruits motor and interoceptive systems, and with Stern’s (2010) vitality affects, which unfold over longer temporal windows and structure the felt continuity of interaction. Together, these perspectives may suggest that therapists dynamically oscillate between rapid simulation-based entrainment and slower interoceptive synchrony in response to relational demands.

The clinical implications follow directly. Mirroring should not be conceived or supervised as a uniform “technique” but as a family of strategies that therapists must flexibly deploy. When client movement is rhythmically coherent and therapist attention is focused on a single interaction partner, sub-second temporal alignment between client and therapist can emerge. In DMT theory, such fine-grained entrainment has been associated with the experience of empathic connection and relational closeness (Tortora, 2009). Under triadic or otherwise high-load conditions, therapists appear to shift strategically toward recurrent structural synchrony, preserving group stability through vertical alignment and postural scaffolding. This echoes clinical reports that embodied attention must be continually redistributed—sometimes being pulled by the more expressive partner, sometimes deliberately directed to stabilise the quieter client to avoid marginalisation (Meekums, 2020). Our findings suggest that, within the constrained timeframe of the triadic task, the therapist’s embodied responses were often drawn toward the most rhythmically and affectively expressive client. While such asymmetries are not inherently problematic, they raise important considerations regarding the distribution of embodied attention within group-based DMT. Crucially, the present analyses necessarily adopt a therapist–client dyadic perspective, reflecting the structure of the motion-capture modelling rather than the full clinical logic of group work. From a clinical standpoint, DMT interventions in group contexts do not aim to validate each client solely through direct therapist mirroring. Rather, therapists work to contain and shape the collective movement field, recognising that clients influence one another’s rhythms, postures, and spatial organisation over time. In this sense, more expressive clients may initially exert greater influence on group rhythm, while the movements of quieter or less expressive clients are often approached more gradually, allowing safety, tolerance, and readiness for exposure to develop. Within short interaction windows, close, fine-grained mirroring of a quieter client may be clinically inappropriate or even intrusive, particularly if it risks overwhelming the individual or destabilising the group field. Interpreted in this light, the present findings do not suggest a failure of equitable attunement but rather illustrate how therapists may temporarily prioritise stabilisation of the group rhythm through more legible movement signals, while supporting less expressive clients via indirect strategies such as spatial anchoring, postural containment, or shared low-frequency movement patterns. Over longer therapeutic timescales, these strategies would typically be complemented by more explicit exploration of quieter clients’ movement material as trust and tolerance increase. Nevertheless, the results highlight an important training and ethical implication: therapists must remain reflexively aware of how expressive salience can bias embodied attention, particularly in early or time-limited group interactions. Training curricula and supervision should therefore emphasise not only sensitivity to micro-timing and resonance, but also intentional modulation of attention across the group, including skills for gradually inviting less expressive clients into shared movement without premature exposure. Tools such as video review or motion-capture–based feedback may offer valuable means of examining tacit embodied habits and supporting reflective practice. Seen in this way, mirroring in group DMT is best understood not as a fixed dyadic exchange, but as a dynamic, distributive practice unfolding over time, in which responsibility lies with the therapist to balance containment, inclusion, and ethical attunement across the evolving group field.

The present results sharpen a central mechanistic question: what occurs in the brain and body when clients experience mirroring as satisfying and transformative? Psychotherapy research has consistently associated in-session synchrony with alliance and outcome (Ramseyer & Tschacher, 2011; Tschacher & Meier, 2020), while social neuroscience shows that expressive movement engages premotor, parietal, and insular cortices (Keysers et al., 2010; Decety & Jackson, 2004). Our findings help to specify which neural systems may underpin the distinct forms of mirroring observed. Fine-grained temporal entrainment, exemplified in the female client-therapist dyad, is plausibly subserved by the AON (premotor and inferior parietal cortices) and temporo-parietal circuits that map observed movement into motor codes. By contrast, the triadic shift towards recurrent synchrony may implicate interoceptive–affective systems in anterior insula and anterior cingulate, regions central to sustaining shared bodily states and co-regulation of arousal. Executive control regions, including prefrontal cortex and supplementary motor areas, may arbitrate between these modes under divided attention, biasing therapists toward slower stabilising strategies when micro-timing becomes costly.

A rigorous next step is to time-lock kinematic indices of both entrainment and recurrence to concurrent biomarkers. Dual-EEG or fNIRS hyperscanning could be used to test whether episodes of high behavioural mirroring accuracy or temporal alignment co-occur with inter-brain coherence and oscillatory signatures of sensorimotor coupling (Dumas et al., 2010; Tognoli & Kelso, 2014). Peripheral measures such as heart-rate variability and respiration can index autonomic co-regulation during these same episodes. Interoceptive processes should be central to this programme: heartbeat-evoked potentials, insular activation, and validated interoceptive accuracy/awareness tasks can probe whether recurrent synchrony scaffolds clients’ capacity to sense and regulate internal states—long proposed as a mechanism of change in body-oriented therapies (Craig, 2009; Payne, 2017). By grounding future neuroimaging hypotheses in precisely defined behavioural markers, this research moves beyond phenomenology toward mechanism, demonstrating not only that mirroring is experienced as therapeutic, but that it is underwritten by identifiable neural processes with potential for plastic change across the course of therapy.

A longitudinal perspective is equally critical. If mirroring supports change, one would anticipate pre–post modulation of AON and interoceptive networks after a course of DMT: stronger premotor–parietal connectivity during action-bservation/imitation, enhanced coupling of insula with motor regions during embodied perspective-taking, and EEG evidence of altered mu/beta reactivity to biological motion. These neural shifts should correlate with in-session kinematic measures (e.g., increased phase synchrony or higher jRQA determinism) and with improvements in alliance and symptom measures, thereby triangulating subjective experience, behaviour and biology.

### Limitations

Limitations warrant caution. The idiographic design (one therapist, two clients) enhances ecological validity but constrains generalisability; replication across therapists, client demographics and diagnostic groups is required. The analysed segments (∼40–45 s) were intentionally selected for salient mirroring; future work should embed these analyses in full-session time-courses, examine sensitivity to window length, and include baseline non-mirroring periods. Kinematics were centre-of-mass and participant-centric; subsequent studies should prefer global laboratory frames, ground distances explicitly in SI units with normalisation to stature/initial separation, and incorporate segmental features (hands, head, trunk orientation) to capture clinically meaningful nuances. Method variance remains an issue: machine-learning models can over-leverage low-frequency content unless paired with phase-based and multivariate controls; we therefore treat convergences across synchrony, causality and recurrence as necessary rather than any single metric being sufficient. Finally, we did not include concurrent neural or autonomic measures; given evidence that entrainment and recurrent coupling rely on partially distinct systems (Novembre & Keller, 2014; Novembre et al., 2017; Craig, 2009), future multimodal designs are needed to test mechanism and track plasticity.

### Conclusions

In conclusion, therapist mirroring in DMT is context-sensitive, comprising distinct yet complementary forms of embodied coordination. Dyads in which client movement exhibits a coherent and rhythmically structured expressive profile tended to show stronger fine-grained temporal entrainment and clearer directional predictability, whereas less temporally structured movement was associated with reduced entrainment and greater reliance on spatial or recurrent forms of coordination. In triadic interactions, temporal micro-alignment was further attenuated, with coordination instead characterised by more stable, low-frequency structural synchrony supporting postural organisation. Situated against DMT practice, developmental accounts of affect attunement, and contemporary models of synchrony and embodied simulation, these results motivate a translational research programme integrating kinematics, hyperscanning, autonomic physiology, and longitudinal imaging. By linking subjective reports, behavioural synchrony measures, and indices of neural and interoceptive coupling, future work could test whether mirroring episodes characterised by high attunement are not only experienced as salient within therapy but also correspond to measurable changes in brain–body coordination.

### Ethics Statement

This study was conducted in accordance with the Declaration of Helsinki and approved by the Institutional Ethics Committee of Edge Hill University. All participants provided written informed consent prior to participation, with assurances regarding anonymity, confidentiality, and the voluntary nature of their involvement.

## Data Availability Statement

The anonymised datasets and analysis scripts supporting the conclusions of this article are available on the Open Science Framework (OSF) via a private, anonymised reviewer link: https://osf.io/sudwj/overview?view_only=d85c21d9a93d45a5adb43c69a6acca58

## Acknowledgements

The authors acknowledge that the conceptual development and interdisciplinary discussions informing this study were supported by an Arts and Humanities Research Council (AHRC) Research Network award to V.C. and V. K. The authors also thank the three participants for their time, engagement, and willingness to take part in the study.

## References

1. ADMP UK. (2025). The Association for Dance Movement Psychotherapy UK. https://admp.org.uk/

2. Aithal, S., Moula, Z., Karkou, V., Karaminis, T., Powell, J., & Makris, S. (2021). A systematic review of the contribution of dance movement psychotherapy to the well-being of children with autism spectrum disorders. Frontiers in Psychology, 12, 719673. 10.3389/fpsyg.2021.719673

3. Barsalou, L. W. (2008). Grounded cognition. Annual Review of Psychology, 59, 617–645. 10.1146/annurev.psych.59.103006.093639

4. Beebe, B., & Lachmann, F. M. (2002). Infant research and adult treatment: Co-constructing interactions. Analytic Press.

5. Bradt, J., Shim, M., & Goodill, S. W. (2015). Dance/movement therapy for improving psychological and physical outcomes in cancer patients. Cochrane Database of Systematic Reviews, (1), CD007103. 10.1002/14651858.CD007103.pub3

6. Cascio, C. J., Moore, D., & McGlone, F. (2019). Social touch and human development. Developmental Cognitive Neuroscience, 35, 5–11. 10.1016/j.dcn.2018.04.009

7. Cheng, Y., Lee, P. L., Yang, C. Y., Lin, C. P., Hung, D., & Decety, J. (2010). Gender differences in the mu rhythm of the human mirror-neuron system. PLoS ONE, 5(5), e2113. 10.1371/journal.pone.0002113

8. Christov-Moore, L., Simpson, E. A., Coudé, G., Grigaityte, K., Iacoboni, M., & Ferrari, P. F. (2014). Empathy: Gender effects in brain and behavior. Neuroscience & Biobehavioral Reviews, 46, 604–627. 10.1016/j.neubiorev.2014.09.001

9. Craig, A. D. (2009). How do you feel—now? The anterior insula and human awareness. Nature Reviews Neuroscience, 10(1), 59–70. 10.1038/nrn2555

10. Decety, J., & Jackson, P. L. (2004). The functional architecture of human empathy. Behavioral and Cognitive Neuroscience Reviews, 3(2), 71–100. 10.1177/1534582304267187

11. De Jaegher, H., & Di Paolo, E. (2007). Participatory sense-making: An enactive approach to social cognition. Phenomenology and the Cognitive Sciences, 6(4), 485–507. 10.1007/s11097-007-9076-9

12. de Witte, M., Bradt, J., Aithal, S., Flynn, L., Karkou, V., Koch, S. C., Orkibi, H., Sajnani, N., Berberian, M., Fietje, N., Miranda, J., Baker, F., & Lampit, A. (2025). The effects of arts-based interventions in the treatment and management of non-communicable diseases: An umbrella review and meta-analyses (Version 1) [Preprint]. Research Square. 10.21203/rs.3.rs-5961850/v1

13. de Witte, M., Orkibi, H., Zarate, R., Karkou, V., Sajnani, N., Malhotra, B., & Koch, S. C. (2021). From therapeutic factors to mechanisms of change in the creative arts therapies: A scoping review. Frontiers in Psychology, 12, 678397. 10.3389/fpsyg.2021.678397

14. Dumas, G., Nadel, J., Soussignan, R., Martinerie, J., & Garnero, L. (2010). Inter-brain synchronization during social interaction. PLoS ONE, 5(8), e12166. 10.1371/journal.pone.0012166

15. Fischman, D. (2009). Therapeutic relationships and kinesthetic empathy. In S. Chaiklin & H. Wengrower (Eds.), The art and science of dance/movement therapy: Life is dance (pp. 33–53). Routledge.

16. Fischer, A. H., & LaFrance, M. (2015). What drives the smile and the tear: Why women are more emotionally expressive than men. Emotion Review, 7(1), 22–29. 10.1177/1754073914544406

17. Fonagy, P., & Allison, E. (2014). The role of mentalizing and epistemic trust in the therapeutic relationship. Psychotherapy, 51(3), 372–380. 10.1037/a0036505

18. Foster, S. L. (2011). Choreographing empathy: Kinesthesia in performance. Routledge.

19. Gallagher, S. (2005). How the body shapes the mind. Oxford University Press.

20. Gallese, V. (2003). The manifold nature of interpersonal relations: The quest for a common mechanism. Philosophical Transactions of the Royal Society B: Biological Sciences, 358(1431), 517–528. 10.1098/rstb.2002.1234

21. Gallese, V. (2009). Mirror neurons, embodied simulation, and the neural basis of social identification. Psychoanalytic Dialogues, 19(5), 519–536. 10.1080/10481880903231910

22. Gallese, V., & Sinigaglia, C. (2011). What is so special about embodied simulation? Trends in Cognitive Sciences, 15(11), 512–519. 10.1016/j.tics.2011.09.003

23. Hove, M. J., & Risen, J. L. (2009). It’s all in the timing: Interpersonal synchrony increases affiliation. Social Cognition, 27(6), 949–960. 10.1521/soco.2009.27.6.949

24. Karkou, V., & Sanderson, P. (2006). Arts therapies: A research-based map of the field. Elsevier.

25. Karkou, V., Aithal, S., Zubala, A., & Meekums, B. (2019). Effectiveness of dance movement therapy in the treatment of adults with depression: A systematic review with meta-analyses. Frontiers in Psychology, 10, 936. 10.3389/fpsyg.2019.00936

26. Karkou, V., Aithal, S., Richards, M., Hiley, E., & Meekums, B. (2023). Dance movement therapy for dementia. Cochrane Database of Systematic Reviews, (8), CD011022. 10.1002/14651858.CD011022.pub3

27. Kattenstroth, J. C., Kalisch, T., Holt, S., Tegenthoff, M., & Dinse, H. R. (2010). Six months of dance intervention enhances postural, sensorimotor, and cognitive performance in elderly adults. Frontiers in Aging Neuroscience, 2, 31. 10.3389/fnagi.2010.00031

28. Koch, S. C. (2014). Embodied affectivity: On moving and being moved. Frontiers in Psychology, 5, 508. 10.3389/fpsyg.2014.00508

29. Koch, S. C., & Fischman, D. (2011). Embodied enactive dance/movement therapy. American Journal of Dance Therapy, 33(1), 57–72. 10.1007/s10465-011-9105-4

30. Koch, S. C., Kunz, T., Lykou, S., & Cruz, R. (2014). Effects of dance movement therapy and dance on health-related psychological outcomes: A meta-analysis. The Arts in Psychotherapy, 41(1), 46–64. 10.1016/j.aip.2013.10.004

31. Koch, S. C., Riege, R. F., Tisborn, K., Biondo, J., Martin, L., & Beelmann, A. (2019). Effects of dance movement therapy and dance on health-related psychological outcomes: A meta-analysis update. Frontiers in Psychology, 10, 1806. 10.3389/fpsyg.2019.01806

32. Koole, S. L., & Tschacher, W. (2016). Synchrony in psychotherapy: A review and an integrative framework for the therapeutic alliance. Frontiers in Psychology, 7, 862. 10.3389/fpsyg.2016.00862

33. Leichsenring, F., Steinert, C., Rabung, S., & Ioannidis, J. P. A. (2022). The efficacy of psychotherapies and pharmacotherapies for mental disorders in adults: An umbrella review. World Psychiatry, 21(1), 133–145. 10.1002/wps.20941

34. Lyons, S., Karkou, V., Roe, B., Meekums, B., & Richards, M. (2018). Evidence for dance movement therapy in older adults with dementia. The Arts in Psychotherapy, 60, 32–40. 10.1016/j.aip.2018.03.006

35. Meekums, B. (2002). Dance movement therapy: A creative psychotherapeutic approach. SAGE. 10.4135/9781446217986

36. Meekums, B. (2020). Dance movement therapy for trauma survivors: A qualitative pilot study. The Arts in Psychotherapy, 69, 101664. 10.1016/j.aip.2020.101664

37. Meekums, B., Karkou, V., & Nelson, E. A. (2015). Dance movement therapy for depression. Cochrane Database of Systematic Reviews, (2), CD009895. 10.1002/14651858.CD009895.pub2

38. Moula, Z., Aithal, S., Karkou, V., & Powell, J. (2020). Child-focused outcomes of arts therapies delivered in primary schools: A systematic review. Children and Youth Services Review, 112, 104928. 10.1016/j.childyouth.2020.104928

39. Moula, Z., Powell, J., Brocklehurst, S., & Karkou, V. (2022). School-based dance movement psychotherapy for children with emotional and behavioural difficulties. Frontiers in Psychology, 13, 883334. 10.3389/fpsyg.2022.883334

40. Payne, H. (Ed.). (1992). Dance movement therapy: Theory and practice. Routledge.

41. Payne, H. (2017). Essentials of dance movement psychotherapy. Routledge.

42. Ren, J., & Xia, J. (2013). Dance therapy for schizophrenia. Cochrane Database of Systematic Reviews, (10), CD006868. 10.1002/14651858.CD006868.pub3

43. Rogers, C. R. (1951/1992). Client-centred therapy. Constable.

44. Roth, A. D., & Fonagy, P. (2006). What works for whom: A critical review of psychotherapy research. Guilford Press.

45. Zhang, Q., Wang, J., & Neitzel, A. (2023). School-based mental health interventions targeting depression or anxiety: A meta-analysis. Journal of Youth and Adolescence, 52, 195–217. 10.1007/s10964-022-01684-4

46. Zubala, A., & Karkou, V. (2015). Dance movement psychotherapy practice in the UK. Body, Movement and Dance in Psychotherapy, 10(1), 21–38. 10.1080/17432979.2014.920918

