## Supplementary material for "Tracing the invisible: Quantifying mirroring and embodied attunement in dyadic and triadic Dance Movement Therapy"

**Supplementary Materials**

**S1. Motion-capture preprocessing and data preparation**

**S1.1 Segment selection**

Analyses focused on short segments extracted from full Dance Movement Therapy sessions under both dyadic and triadic conditions. Segments corresponded to periods in which the therapist explicitly engaged in mirroring, as defined by the intervention protocol and independently validated, and reflected sustained participation by all parties during the therapist-led mirroring phase. Each analysed segment lasted approximately 40–45 seconds, providing a standardised temporal window for assessing interpersonal coordination.

**S1.2 Reconstruction and labelling (Qualisys Track Manager)**

Marker trajectories were reconstructed and labelled in Qualisys Track Manager (QTM; version 2024.3). Short gaps of up to 10 consecutive frames were interpolated using QTM’s built-in polynomial gap-filling procedure. Labelled trials were cropped to the time window of interest and exported for downstream processing.

**S1.3 Downstream processing and analysis environment**

Visual3D (C-Motion, USA) was used exclusively for biomechanical modelling and extraction of trunk centre-of-mass trajectories. All subsequent analyses, statistical modelling, and figure generation were performed using custom R and Python scripts.

**S1.4 Filtering**

Marker/CoM trajectories were low-pass filtered using a fourth-order, zero-lag Butterworth filter with a 6 Hz cut-off frequency.

**S1.5 Coordinate conventions and axes**

Whole-body movement was represented via the centre of mass (CoM) of the trunk segment, treated as a proxy for global body motion. CoM displacement was analysed along the three principal axes:

- ML (medio–lateral; side-to-side)
- AP (antero–posterior; forwards–backwards)
- V (vertical; up–down)

These axis conventions are consistent across dyadic and triadic analyses.

**S1.6 CoM extraction, normalisation, and distance computation**

For each participant and trial, three-dimensional trunk CoM trajectories were extracted and analysed along ML, AP, and V axes. Where required by specific analyses, CoM trajectories were normalised within trials to facilitate comparability while preserving temporal structure.

To characterise spatial organisation, three-dimensional Euclidean distance between therapist and client trunk CoM trajectories was computed and expressed relative to the initial separation (normalised inter-CoM distance).

**S1.7 Standardisation for time-series analyses**

Where required (cross-correlation, synchronisation, predictive modelling, Granger causality, VAR), CoM time series were z-standardised within trials to control for amplitude differences while preserving the temporal structure of movement dynamics.

**S2. Dyadic Mirroring with the Female Client**

This section reports the full set of dyadic analyses for the female client–therapist interaction that were summarised in the main text. All analyses were conducted on z-standardised centre-of-mass (CoM) trajectories and are reported here in full to preserve axis-specific statistics and directional detail.

**S2.1 Cross-Correlation Analysis**

Cross-correlation functions were computed separately for the medio–lateral (ML), antero–posterior (AP), and vertical (V) axes using a ±5 s lag window.

Along the ML axis, the maximum positive cross-correlation was weak (r = .137) and occurred at a lag of +3.00 s, indicating delayed coupling. In contrast, the strongest negative correlation was substantially larger in magnitude (r = −.706) and occurred at −0.34 s, indicating a pronounced anti-phase relationship at a sub-second delay. This pattern suggests that lateral displacements of the therapist were temporally coupled to those of the client but occurred in the opposite direction, consistent with left–right mirroring dynamics.

In the AP axis, cross-correlation peaked at r = .749 at a lag of −0.73 s, indicating that the client’s forward–backward movements consistently preceded those of the therapist by less than one second.

The V axis showed the strongest temporal alignment, with a peak correlation of r = .977 at −0.39 s, reflecting highly consistent vertical mirroring within a narrow temporal window.

At zero lag, statistically significant concurrent associations were observed in both the AP axis (r = .715, p = .007) and the V axis (r = .967, p = .001). In contrast, the ML axis exhibited a significant negative concurrent association (r = −.628, p = .017), consistent with the anti-phase lateral dynamics identified in the lagged analysis.

Together, these results indicate strong and temporally precise mirroring in the vertical and antero–posterior dimensions, alongside structured anti-phase coordination in the medio–lateral axis.

**S2.2 Phase Synchronisation Analysis**

Phase synchronisation was quantified using the Synchronisation Index (SI) across the 0.1–3 Hz frequency range, with statistical significance assessed via permutation testing.

In the ML axis, synchronisation was low (SI = 0.338) and did not exceed the permutation-derived significance threshold (p = .475). Similarly, synchronisation in the AP axis was low (SI = 0.292) and non-significant (p = .305).

By contrast, the V axis exhibited statistically significant phase synchronisation (SI = 0.626, p = .032), exceeding the 95th percentile of the null distribution. This indicates reliable phase-locking between therapist and client exclusively in the vertical dimension.

These findings demonstrate that, despite lagged temporal coupling in multiple axes, statistically significant phase-based synchrony emerged only for vertical movement in the female dyad.

**S2.3 Spatial Organisation**

Spatial organisation was assessed using normalised inter-centre-of-mass (CoM) distance, expressed relative to the initial therapist–client separation (see also Figure S1).

The mean normalised inter-CoM distance was 1.042 (SD = 0.416), with observed values ranging from 0.238 to 2.306. These values indicate that, on average, the therapist and client maintained a spatial separation close to their initial configuration, while exhibiting substantial fluctuations over time.

The observed variability reflects dynamic modulation of interpersonal spacing during the mirroring task rather than maintenance of a fixed spatial configuration.

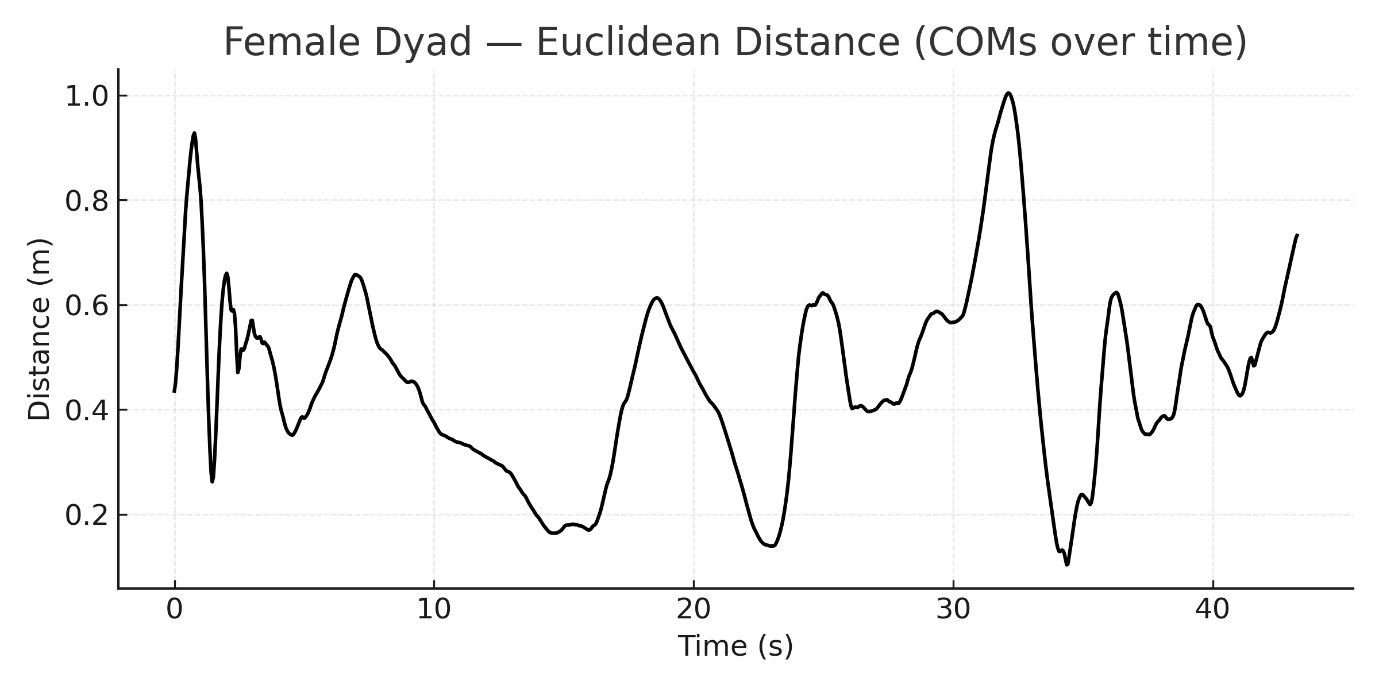

***Figure S1.*** *Female dyad — Euclidean distance between centres of mass (COM). The plot shows the three-dimensional Euclidean distance in metres between the therapist and the female client across the duration of the trial. The curve represents the absolute inter-COM distance, while the legend identifies the trajectory as Euclidean Distance (m). This measure reflects the spatial proximity of the dyad, with smaller values indicating closer positioning and larger values indicating greater separation.*

**S2.4 Random Forest Predictive Coupling**

Directional predictive coupling was examined using Random Forest regression models with three-block time-ordered cross-validation (see also Figure S2).

In the ML axis, client-to-therapist prediction was extremely strong (R² = .984, MSE = 0.016), whereas therapist-to-client prediction was markedly weaker (R² = .792, MSE = 0.208).

A similar directional asymmetry was observed in the AP axis, where client-to-therapist prediction reached R² = .957 (MSE = 0.042), compared with R² = .860 (MSE = 0.195) for therapist-to-client prediction.

In contrast, predictive accuracy in the V axis was low and comparable in both directions (client → therapist R² ≈ .06, MSE ≈ 0.88; therapist → client R² ≈ .06, MSE ≈ 0.88), indicating limited directional predictability in vertical movement dynamics.

Overall, Random Forest modelling revealed a clear client-driven predictive asymmetry in the horizontal planes, with weak and reciprocal predictability in the vertical dimension.

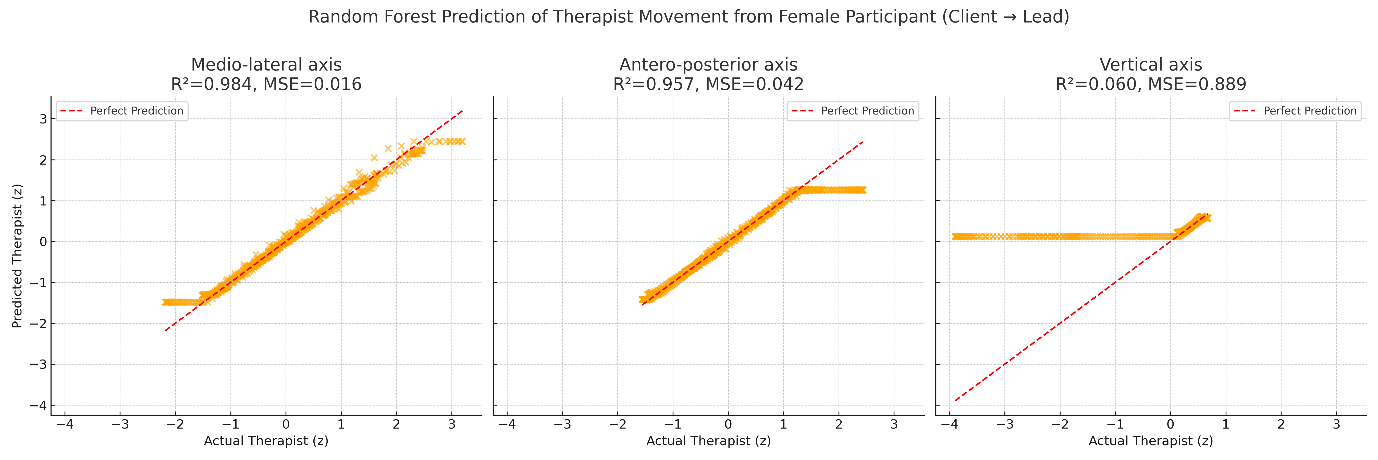

***Figure S2.*** *Random Forest prediction of therapist movement from female client across axes (client → lead). Predicted versus actual therapist movement is shown for the medio-lateral (ML), antero-posterior (AP), and vertical (V) axes. Each panel displays out-of-sample predictions from a Random Forest model trained on client history using eight temporal lags. The dashed red line indicates the line of perfect prediction (y = x). Model performance is reported in each panel as explained variance (R²) and mean squared error (MSE). Prediction accuracy was high for the ML (R² = 0.984, MSE = 0.016) and AP (R² = 0.957, MSE = 0.042) axes, but negligible for the V axis (R² = 0.060, MSE = 0.889).*

**S2.5 Granger Causality Analysis**

Granger causality analyses were conducted on z-standardised CoM trajectories, with lag order selected using the Akaike Information Criterion (maximum of eight lags). All reported effects survived false discovery rate (FDR) correction within condition unless otherwise stated.

In the ML axis, significant bidirectional Granger causality was observed. The client’s past movements significantly predicted the therapist’s future dynamics (F(7, 988) = 6.62, p < .001), while the therapist-to-client pathway was also significant but weaker (F(7, 988) = 2.14, p = .038).

In the AP axis, a pronounced directional asymmetry emerged. Client-to-therapist Granger causality was significant (F(6, 991) = 3.34, p = .003), whereas therapist-to-client causality was not (F(8, 985) = 0.74, p = .655).

In the V axis, strong bidirectional coupling was evident. The client significantly predicted the therapist (F(7, 988) = 13.95, p < .001), and the therapist also exerted a significant influence on the client (F(6, 991) = 3.09, p = .005).

Taken together, Granger causality analyses indicate predominant client-to-therapist directional influence across all movement dimensions, with additional reciprocal coupling in the ML and vertical axes.

**S3. Dyadic Mirroring with the Male Client**

This section reports the complete set of dyadic analyses for the male client–therapist interaction. These results were moved from the main Results section to the Supplementary Materials to preserve space and focus, but are reproduced here in full, including all statistical details.

**S3.1 Cross-Correlation Analysis**

Cross-correlation functions were computed between therapist and male client centre-of-mass (CoM) trajectories for the medio–lateral (ML), antero–posterior (AP), and vertical (V) axes using a ±5 s lag window (see also Figure S3).

Along the ML axis, the cross-correlation function peaked at r = .217 with an optimal lag of +1.89 s, indicating weak client-leading dynamics. Negative correlations were also observed, with a negative peak of r = −.172, suggesting intermittent anti-phase relationships in lateral displacement.

In contrast, the AP axis exhibited stronger coupling. The peak positive correlation reached r = .515 at a lag of −2.10 s, indicating therapist-leading coordination with a strong association according to predefined correlation magnitude benchmarks.

Similarly, the V axis showed a peak correlation of r = .511 at a lag of −1.97 s, again indicating therapist-leading dynamics and strong temporal coupling in vertical movement.

Negative cross-correlation peaks were also evident in the vertical axis (r = −.249), suggesting transient anti-phase relations alongside periods of in-phase coordination.

Taken together, these results indicate that, in the male dyad, temporal synchrony was weak and delayed in the ML dimension but substantially stronger in the AP and V axes, where therapist-led coordination predominated.

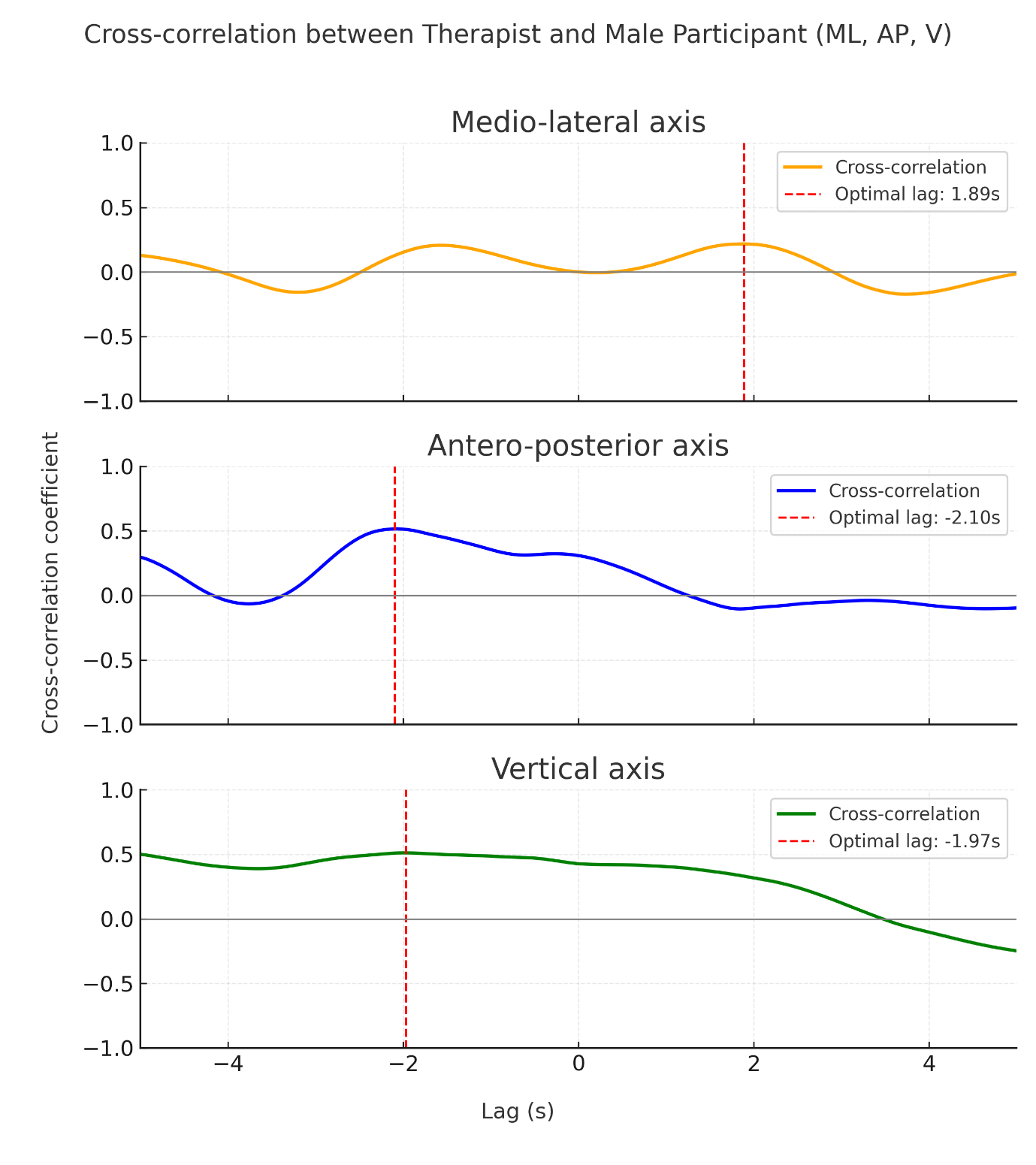

***Figure S3.*** *Cross-correlation between therapist and male client across axes. Each panel shows the cross-correlation function for the medio-lateral (ML), antero-posterior (AP), and vertical (V) axes, with the optimal lag indicated by a dashed red line. Axes are standardised to ±5 s on the x-axis and −1 to 1 on the y-axis.*

**S3.2 Phase Synchronisation Analysis**

Phase-based synchronisation between therapist and male client movements was quantified using the Synchronisation Index (SI), with statistical significance assessed via permutation testing (see also Figure S4).

Synchronisation in the ML axis was very low (SI = 0.076) and did not exceed the permutation-derived significance threshold (p = .891). Similarly, synchronisation in the AP axis was low (SI = 0.145, p = .741).

The V axis showed comparatively higher synchronisation values (SI = 0.257), but this effect also failed to reach statistical significance (p = .541).

These findings indicate that the male dyad did not exhibit statistically significant phase-locking in any movement dimension, in contrast to the significant vertical synchronisation observed in the female dyad.

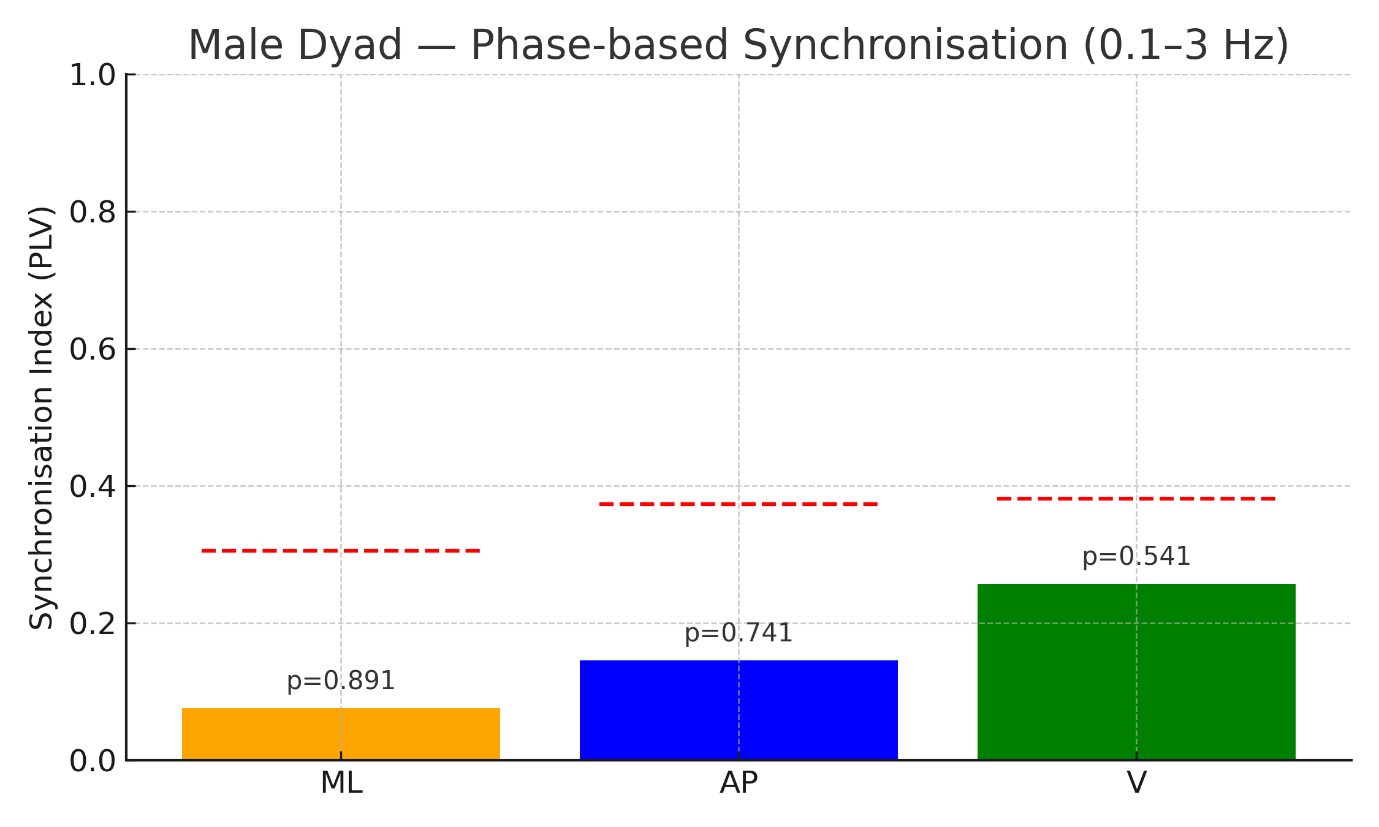

***Figure S4.*** *Phase-based synchronisation for the male dyad. Bars represent the Synchronisation Index (SI) for each axis, with dashed red lines marking the 95th-percentile permutation thresholds. Exact p-values are annotated above the bars.*

**S3.3 Spatial Organisation**

Spatial organisation was assessed using normalised inter-centre-of-mass (CoM) distance, expressed relative to the initial therapist–client separation (see also Figure S5).

The mean normalised inter-CoM distance was 0.762 (SD = 0.362), with observed values ranging from 0.149 to 2.355. These values indicate that, relative to their starting configuration, the therapist and male client frequently reduced their interpersonal spacing, although periods of increased separation were also evident.

The observed variability in normalised distance suggests less stable spatial organisation over the course of the mirroring task compared with the female dyad, reflecting greater fluctuation in interpersonal spacing during the interaction.

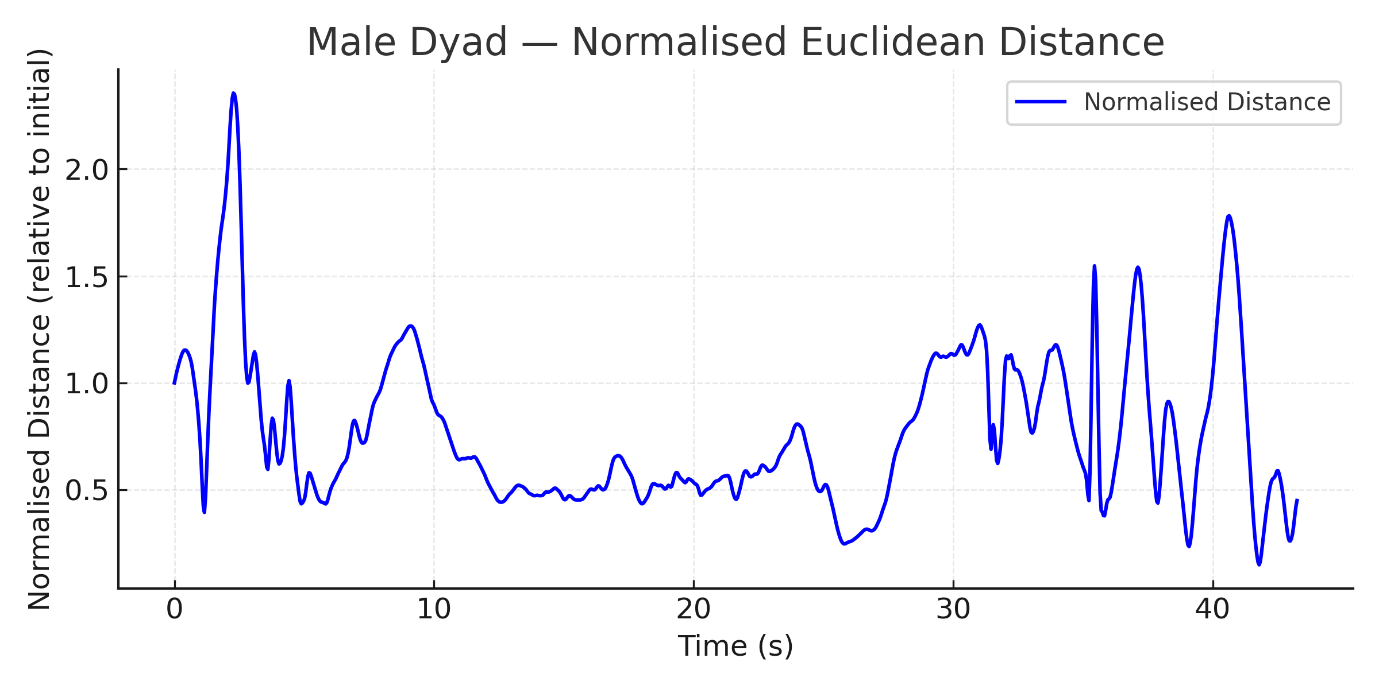

***Figure S5.*** *Euclidean distance between therapist and male client. The graph presents the distance normalised to the dyad’s initial separation, providing a relative index of spacing.*

**S3.4 Random Forest Predictive Coupling**

Directional predictive coupling was examined using Random Forest regression models with three-block time-ordered cross-validation.

Across both the ML and AP axes, the therapist’s movements were more predictive of the client’s subsequent movements than vice versa (see also Figure S6).

In the ML axis, therapist-to-client prediction was strong (R² = .908, MSE = 0.176), whereas client-to-therapist prediction was weaker (R² = .799, MSE = 0.515).

Similarly, in the AP axis, therapist-to-client prediction achieved R² = .921 (MSE = 0.144), exceeding client-to-therapist prediction (R² = .801, MSE = 0.373).

In contrast, the V axis exhibited very high bidirectional predictability, with client-to-therapist prediction yielding R² = .987 (MSE = 0.009) and therapist-to-client prediction R² = .977 (MSE = 0.022). This pattern is consistent with either highly reciprocal coordination or constrained variability in vertical movement within this dyad.

Overall, Random Forest analyses indicate therapist-led predictive coupling in the horizontal planes, alongside tightly coupled or constrained vertical dynamics.

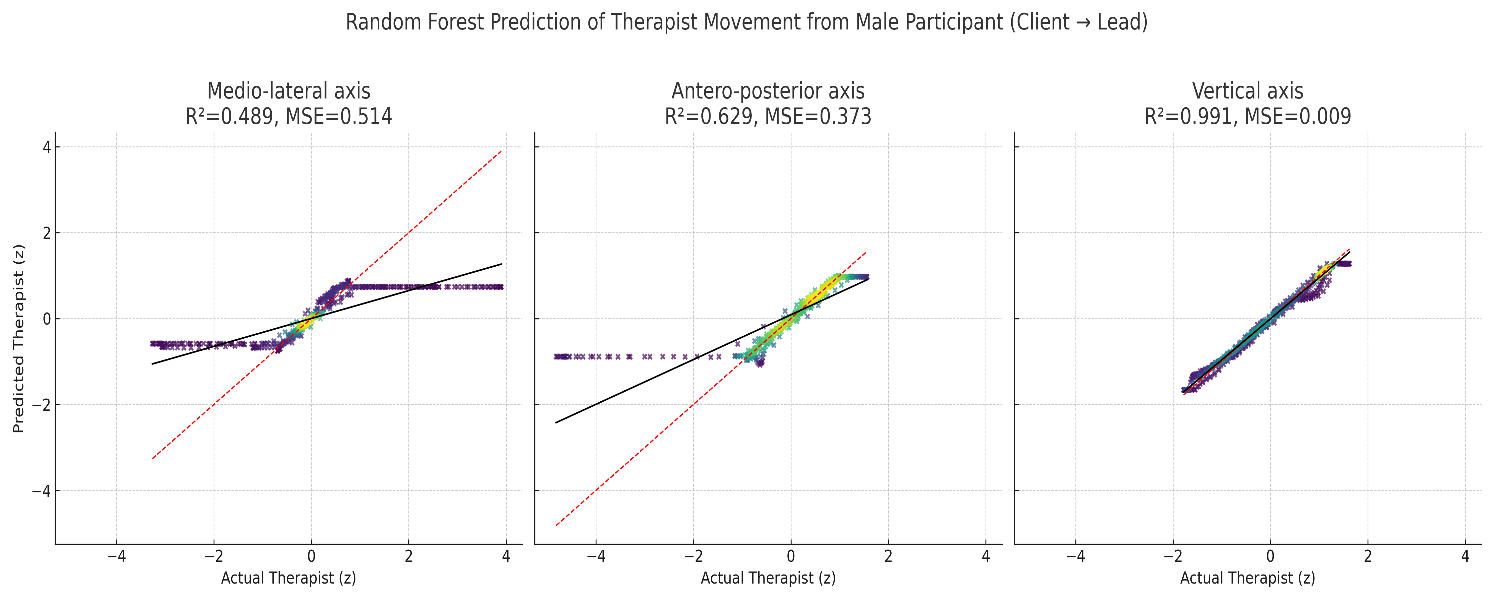

***Figure S6.*** *Random Forest prediction of therapist movement from male client across axes (client → lead). Predicted versus actual therapist movements are shown for the medio-lateral (ML), antero-posterior (AP), and vertical (V) axes. Point density is colour-coded (darker = higher density). The dashed red line indicates perfect prediction (y = x), and the solid black line shows the regression fit. Prediction accuracy was moderate in ML (R² = 0.489, MSE = 0.514) and AP (R² = 0.629, MSE = 0.373), but extremely high in V (R² = 0.991, MSE = 0.009).*

**S3.5 Granger Causality Analysis**

Granger causality analyses were conducted on z-standardised CoM trajectories, with lag order selected using the Akaike Information Criterion (maximum of eight lags). All reported effects survived false discovery rate (FDR) correction within condition unless otherwise stated (see also Figure S7).

Along the ML axis, neither direction showed significant Granger causality (client → therapist: F(6, 991) = 0.41, p = .869; therapist → client: F(8, 985) = 0.63, p = .755).

Similarly, no significant effects were observed in the AP axis (client → therapist: F(8, 985) = 1.45, p = .173; therapist → client: F(8, 985) = 1.05, p = .399).

By contrast, in the V axis there was clear evidence of a unidirectional influence from client to therapist. The client’s past dynamics significantly improved prediction of the therapist’s subsequent movements (F(6, 991) = 3.22, p = .0039), whereas the reverse pathway did not reach significance (F(7, 988) = 0.40, p = .900).

Taken together, these results indicate a selective vertical directional influence from the male client to the therapist, with no reliable Granger causality effects detected in the horizontal movement dimensions.

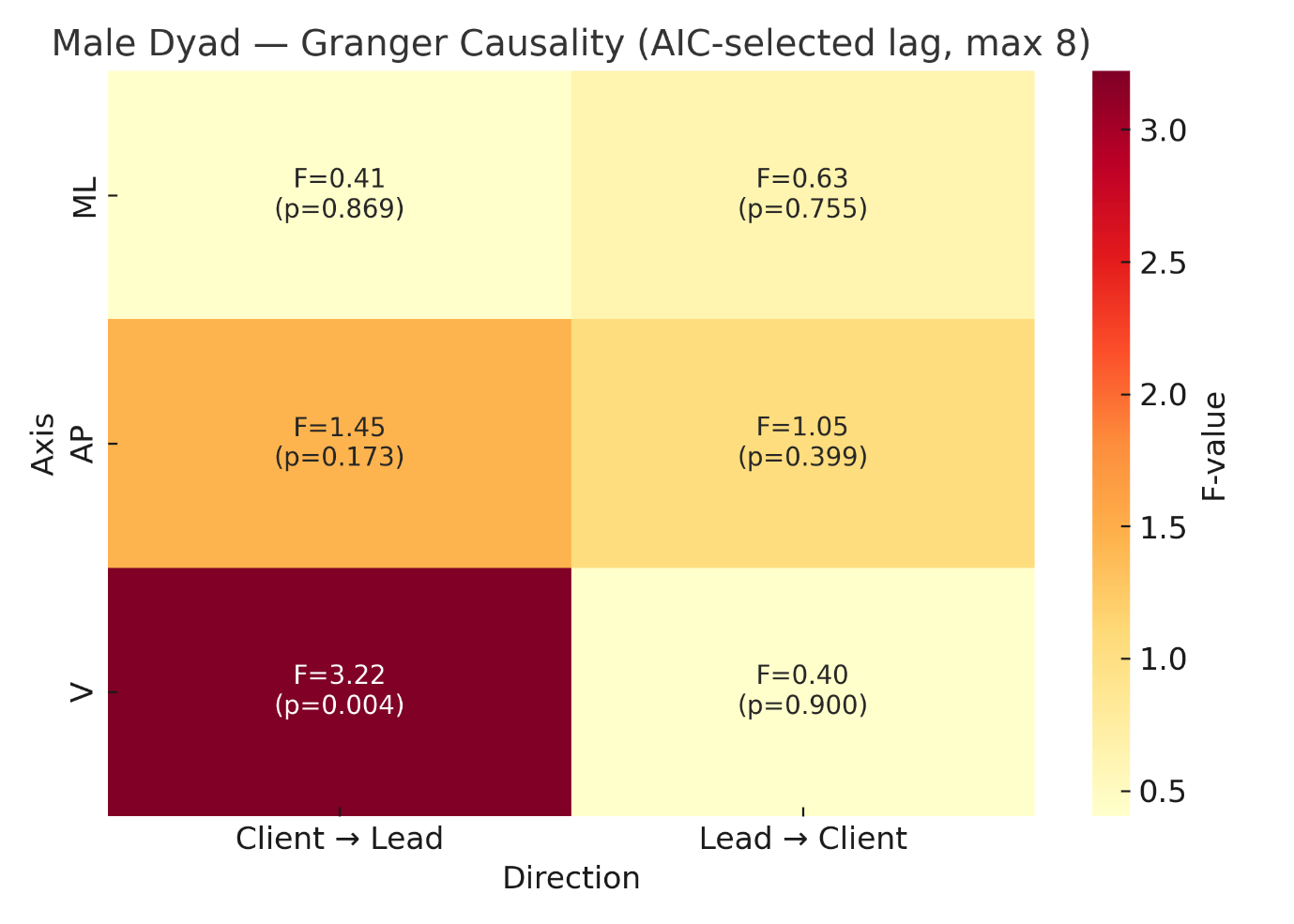

***Figure S7.*** *Granger causality for the male dyad. Heatmap of F-values for Granger causality (AIC-selected lags, max 8) across medio-lateral (ML), antero-posterior (AP), and vertical (V) axes. Columns depict directionality (Client → Lead; Lead → Client). Each cell is annotated with the corresponding F-statistic and p-value. A significant effect is visible for the vertical axis from client to therapist, with no evidence of Granger causality in the other directions/axes.*

**S4. Triadic Mirroring Results**

This section reports the complete results for the triadic mirroring condition, in which the therapist interacted simultaneously with a female and a male client. These analyses were moved from the main Results section to the Supplementary Materials to preserve focus and length but are reproduced here in full, including all statistical details.

**S4.1 Triadic Mirroring with Female Client — Cross-Correlation Analysis**

Cross-correlation functions were computed between therapist and female client centre-of-mass (CoM) trajectories along the medio–lateral (ML), antero–posterior (AP), and vertical (V) axes using a ±5 s lag window (see also Figure S8).

Along the ML axis, associations were weak and temporally delayed. The maximum positive correlation was r = .284 at +5.00 s, while the strongest negative correlation was r = −.362 at +3.60 s, indicating limited alignment and intermittent anti-phase relations.

In the AP axis, coordination was similarly weak and predominantly anti-phasic. The dominant negative peak was r = −.510 at +2.49 s, and no meaningful positive coupling was observed (largest positive value r = −.316 at +3.64 s).

By contrast, the V axis exhibited a strong near-zero-lag association, with a peak correlation of r = .747 at −0.09 s, indicating that the female client’s vertical movements consistently preceded those of the therapist by a fraction of a second.

These results indicate that, within the triadic context, therapist–female client coupling was most pronounced in the vertical dimension, whereas horizontal coordination was weak, delayed, and in part anti-phasic.

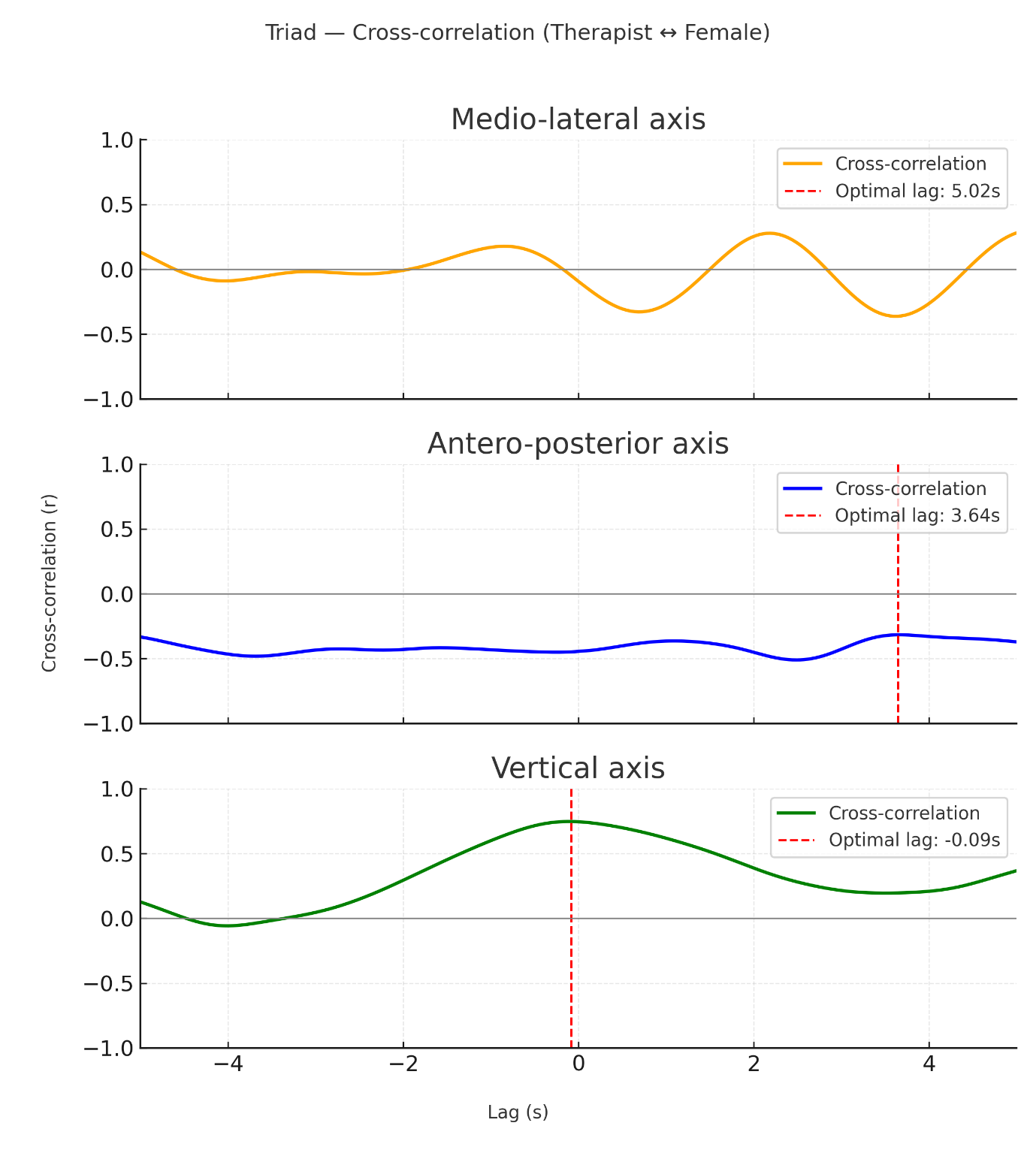

***Figure S8.*** *Cross-correlation between therapist and female client in the triadic task. Each panel shows the correlation function for the ML, AP, and V axes (±5 s window). The dashed red line indicates the lag of peak correlation. Vertical alignment showed strong synchrony at near-zero lag, while ML and AP axes displayed delayed and inconsistent coupling.*

S4.2 Triadic Mirroring with Female Client — Phase Synchronisation Analysis

Phase synchronisation between therapist and female client movements was quantified using the Synchronisation Index (SI), with significance assessed via permutation testing (see also Figure S9).

Synchronisation in the ML axis was low (SI = 0.242) and non-significant (p = .687). Similarly, synchronisation in the AP axis was minimal (SI = 0.035, p = .969).

The V axis showed comparatively higher synchronisation (SI = 0.308), but this effect did not exceed the permutation-derived significance threshold (p = .241).

Thus, unlike the dyadic female interaction, the triadic setting did not support statistically significant phase-locking between therapist and female client in any movement dimension.

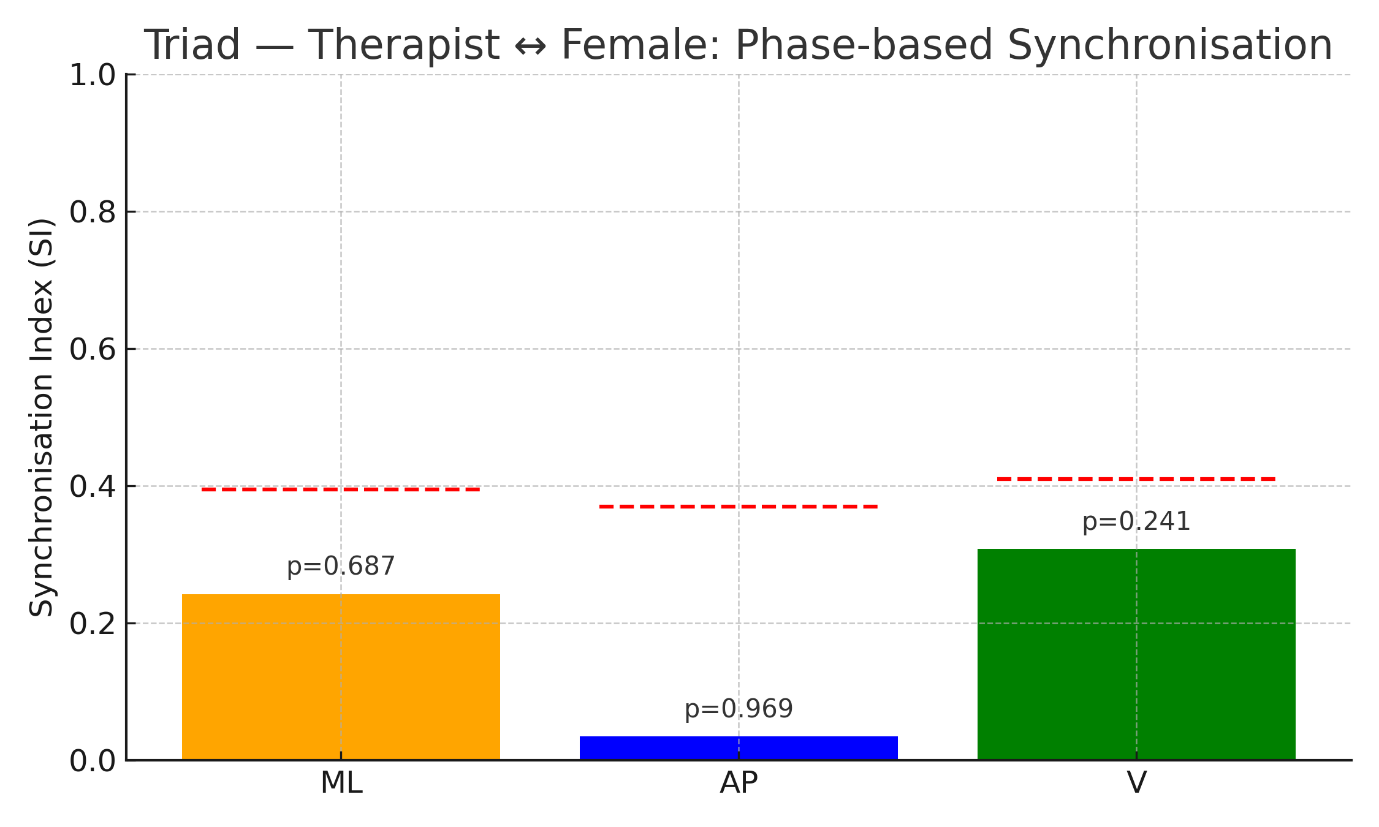

***Figure S9.*** *Synchronisation Index (SI) for therapist–female triadic interaction across ML, AP, and V axes. Bars show observed SI values, dashed red lines mark 95th percentile permutation thresholds, and p-values are annotated.*

**S4.3 Triadic Mirroring with Female Client — Spatial Organisation**

Spatial organisation was assessed using normalised inter-centre-of-mass (CoM) distance, expressed relative to the initial therapist–client separation (see also Figure S10).

The mean normalised inter-CoM distance was 0.809 (SD = 0.278), with values ranging from 0.110 to 1.528. These values indicate that the therapist and female client frequently reduced their interpersonal spacing relative to the starting configuration, while also exhibiting intermittent periods of increased separation.

Overall, the spatial relationship was dynamically modulated rather than fixed throughout the triadic interaction.

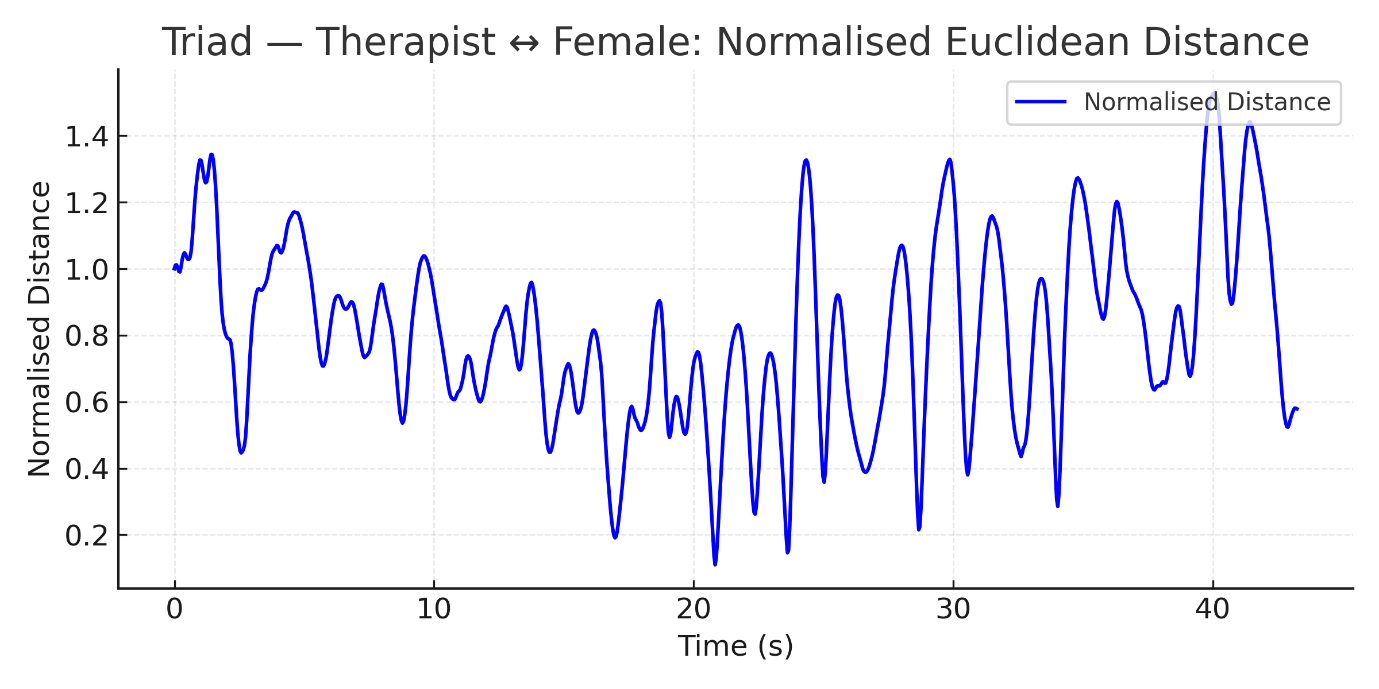

***Figure S10.*** *Euclidean distance between therapist and female client in the triadic task.*

**S4.4 Triadic Mirroring With Female Client — Random Forest Predictive Coupling**

Directional predictive coupling was examined using Random Forest regression models with three-block time-ordered cross-validation (see also Figure S11).

In the ML axis, predictive accuracy was extremely high and nearly identical in both directions (female → therapist: R² = .992, MSE = 0.008; therapist → female: R² = .995, MSE = 0.005).

In the AP axis, a clear directional asymmetry was observed. Female-to-therapist prediction was substantial (R² = .776, MSE = 0.225), whereas therapist-to-female prediction performed poorly (R² = −.263, MSE = 0.175).

In the V axis, predictive accuracy was moderate and broadly symmetric (female → therapist: R² = .734, MSE = 0.268; therapist → female: R² = .659, MSE = 0.703).

These results indicate axis-specific patterns of predictive coupling in the triadic condition, with symmetric predictability in the ML dimension and female-driven predictability in the AP dimension.

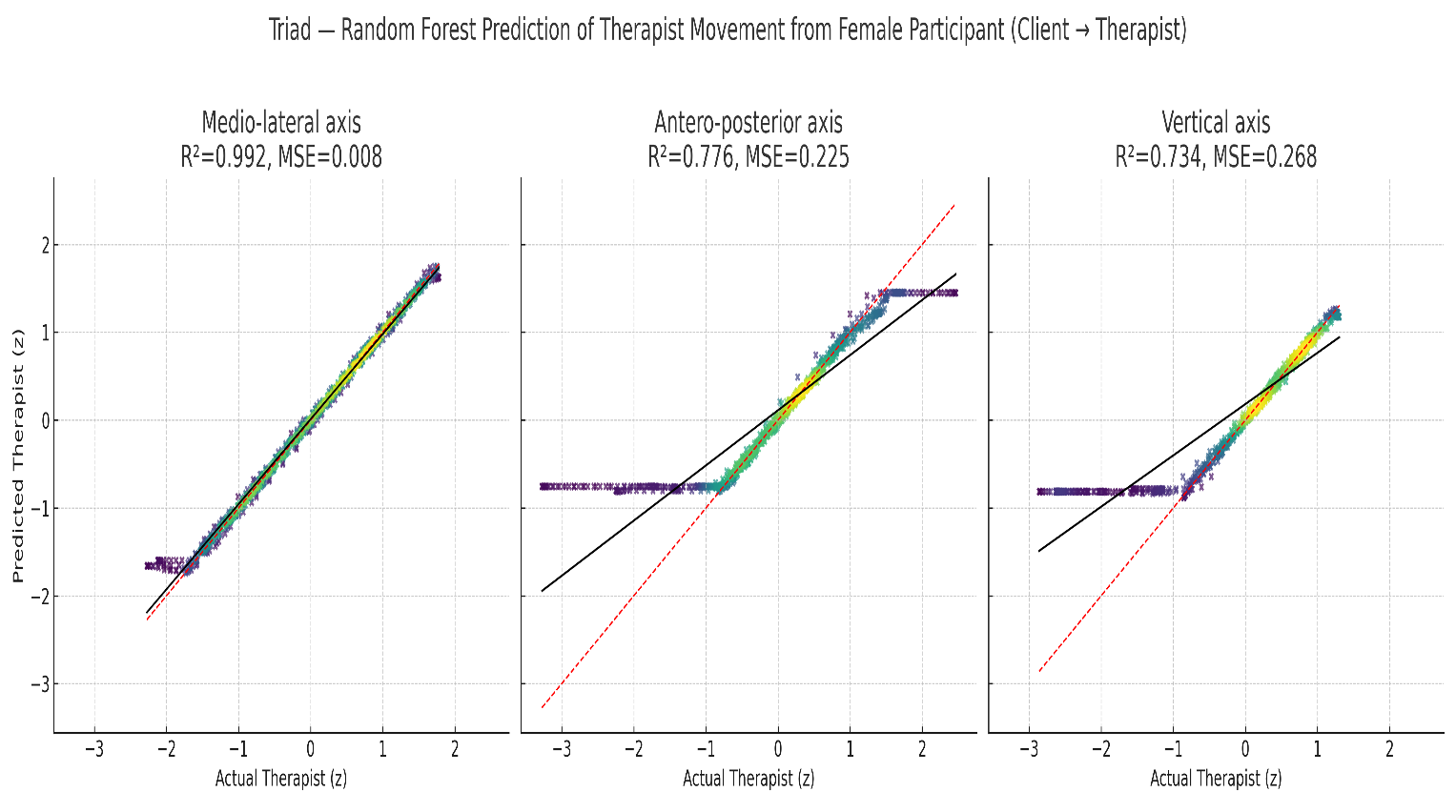

***Figure S11.*** *Density scatter plots of therapist COM trajectories predicted from female client movements in the triadic task. Panels display ML, AP, and V axes with R² and MSE values annotated.*

**S4.5 Triadic Mirroring with Female Client — Granger Causality Analysis**

Granger causality analyses were conducted on z-standardised CoM trajectories, with lag order selected using the Akaike Information Criterion (maximum of eight lags). All reported effects survived false discovery rate (FDR) correction within condition unless otherwise stated (see also Figure S12).

Along the ML axis, neither direction reached statistical significance (female → therapist: F(7, 988) = 1.64, p = .119; therapist → female: F(8, 985) = 1.02, p = .420).

Similarly, no significant effects were observed in the AP axis (female → therapist: F(7, 988) = 0.82, p = .568; therapist → female: F(7, 988) = 0.90, p = .509).

By contrast, the V axis exhibited a significant unidirectional effect from the female client to the therapist (F(7, 988) = 3.48, p = .0011), whereas the reciprocal pathway did not reach significance (F(8, 985) = 1.51, p = .150).

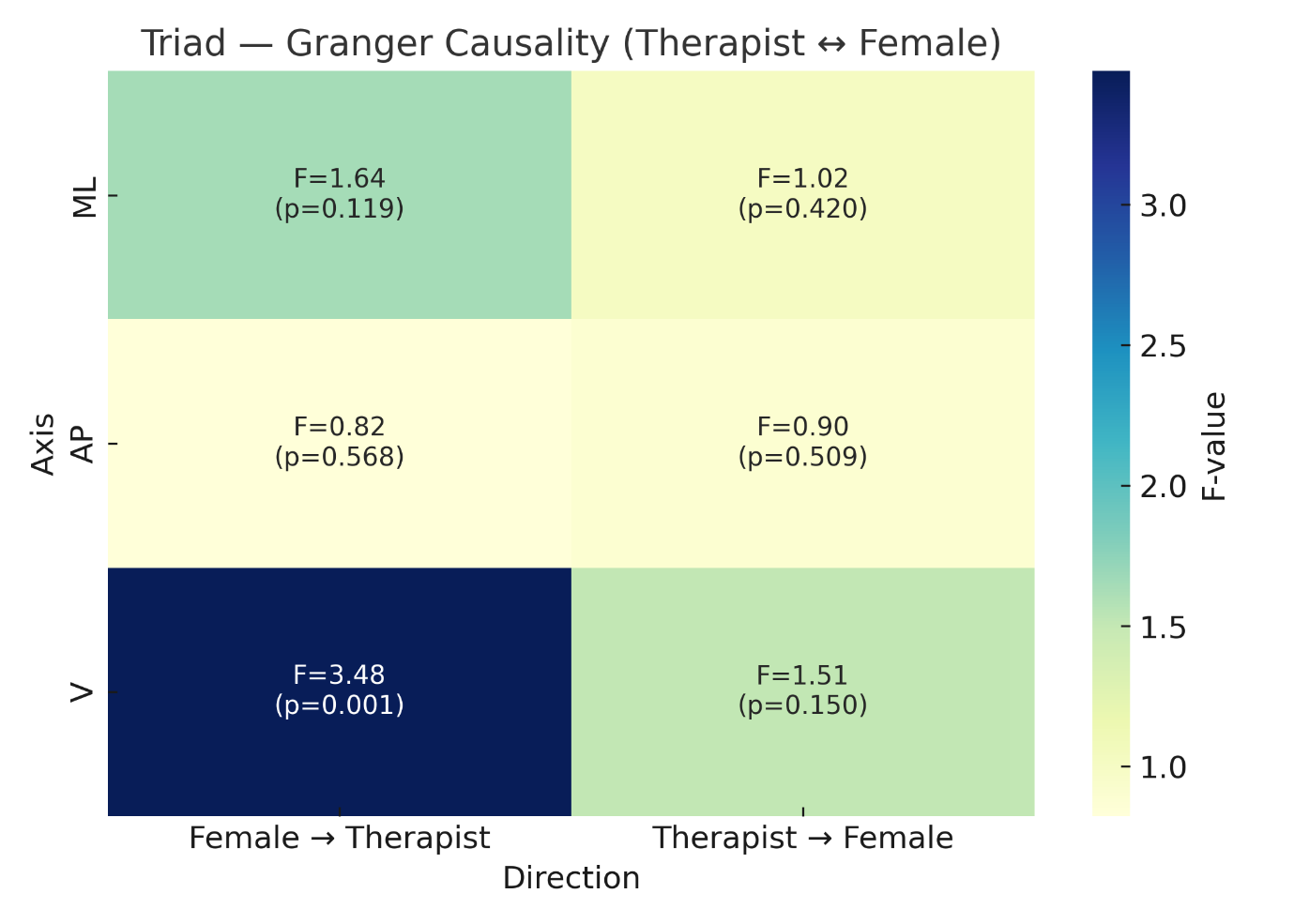

***Figure S12.*** *Granger causality results for therapist ↔ female client in the triadic task. Heatmap shows F-values across ML, AP, and V axes for each direction of influence. Cells are annotated with F-statistics and p-values, with significant causal influence emerging only in the vertical axis from female to therapist.*

**S4.6 Triadic Mirroring with Male Client — Cross-Correlation Analysis**

Cross-correlation functions between therapist and male client CoM trajectories were computed within a ±5 s lag window (see also Figure S13).

In the ML axis, associations were weak and delayed, with a maximum positive correlation of r = .221 at +3.43 s and a negative peak of r = −.413 at +5.00 s.

In the AP axis, no meaningful positive coupling was observed. The cross-correlation function was dominated by negative values, with a peak of r = −.380 at −5.00 s and a near-zero local maximum (r = −.003 at −0.13 s).

By contrast, the V axis exhibited a strong near-zero-lag association (r = .754 at +0.43 s), indicating that therapist vertical movement slightly followed that of the male client.

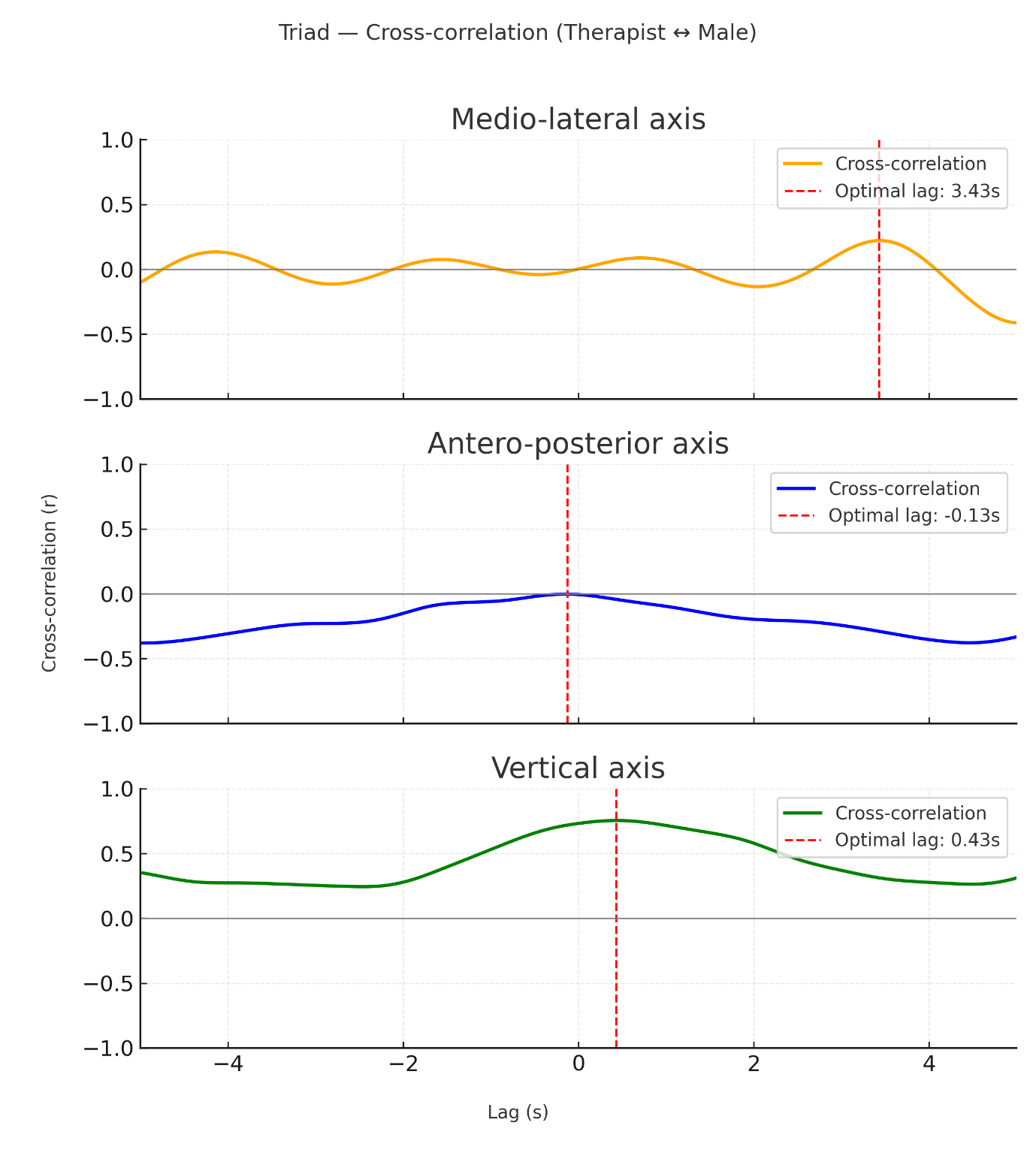

***Figure S13.*** *Cross-correlation between therapist and male client in the triadic task across ML, AP, and V axes (±5 s). The dashed red line marks the lag of the peak positive correlation.*

**S4.7 Triadic Mirroring with Male Client — Phase Synchronisation Analysis**

Phase synchronisation analysis revealed no statistically significant phase-locking between therapist and male client movements in the triadic condition (see also Figure S14).

Synchronisation was very low in the ML axis (SI = 0.050, p = .938). The AP axis showed higher SI values (SI = 0.217), but this effect did not reach significance (p = .453). The V axis exhibited the largest SI values (SI = 0.305), yet synchronisation remained non-significant (p = .508).

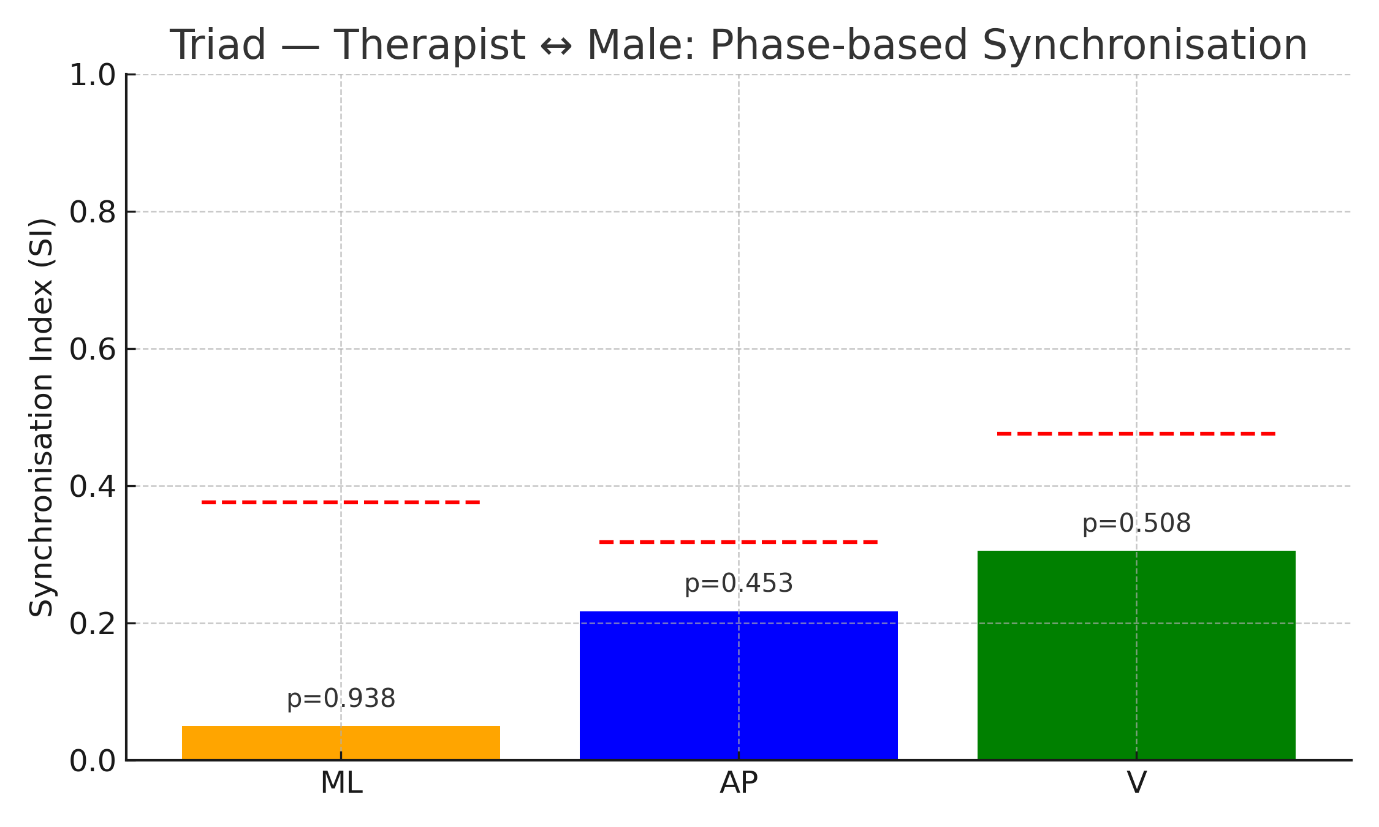

***Figure S14.*** *Synchronisation Index (SI) for therapist–male triadic interaction across ML, AP, and V axes. Bars display observed SI values; dashed red lines indicate 95th-percentile permutation thresholds; p-values are annotated above bars.*

**S4.8 Triadic Mirroring with Male Client — Spatial Organisation**

The mean normalised inter-CoM distance between therapist and male client was 0.753 (SD = 0.309), with values ranging from 0.053 to 1.409 (see also Figure S15).

As in the female triadic interaction, interpersonal spacing was dynamically modulated, with frequent reductions in distance interspersed with periods of increased separation.

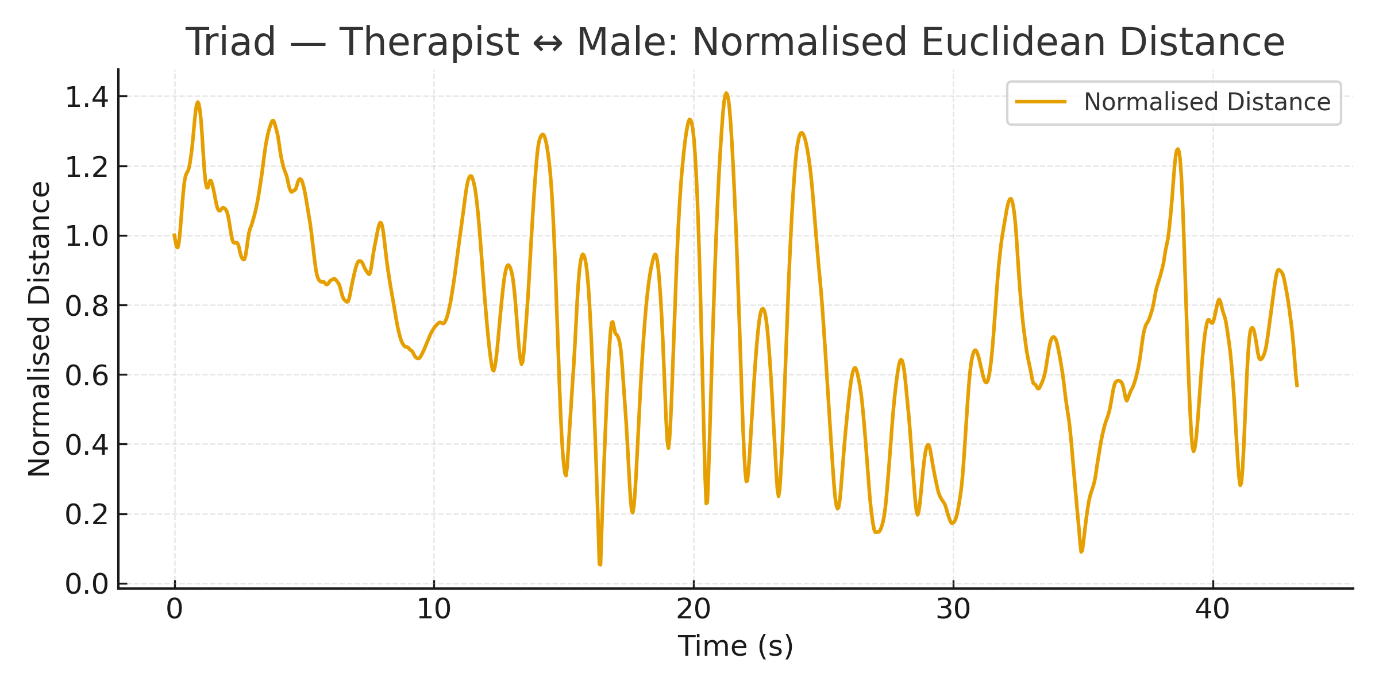

***Figure S15.*** *Euclidean distance between therapist and male client in the triadic task.*

**S4.9 Triadic Mirroring with Male Client — Random Forest Predictive Coupling**

Random Forest analyses revealed axis-specific predictive asymmetries (see also Figure S16).

In the ML axis, predictive accuracy was extremely high and symmetric (male → therapist: R² = .992, MSE = 0.008; therapist → male: R² = .991, MSE = 0.009).

In the AP axis, predictive accuracy favoured male-to-therapist prediction (R² = .779, MSE = 0.222) over therapist-to-male prediction (R² = .706, MSE = 0.291).

In the V axis, predictive accuracy was lower overall but again asymmetric (male → therapist: R² = .716, MSE = 0.286; therapist → male: R² = .592, MSE = 0.409).

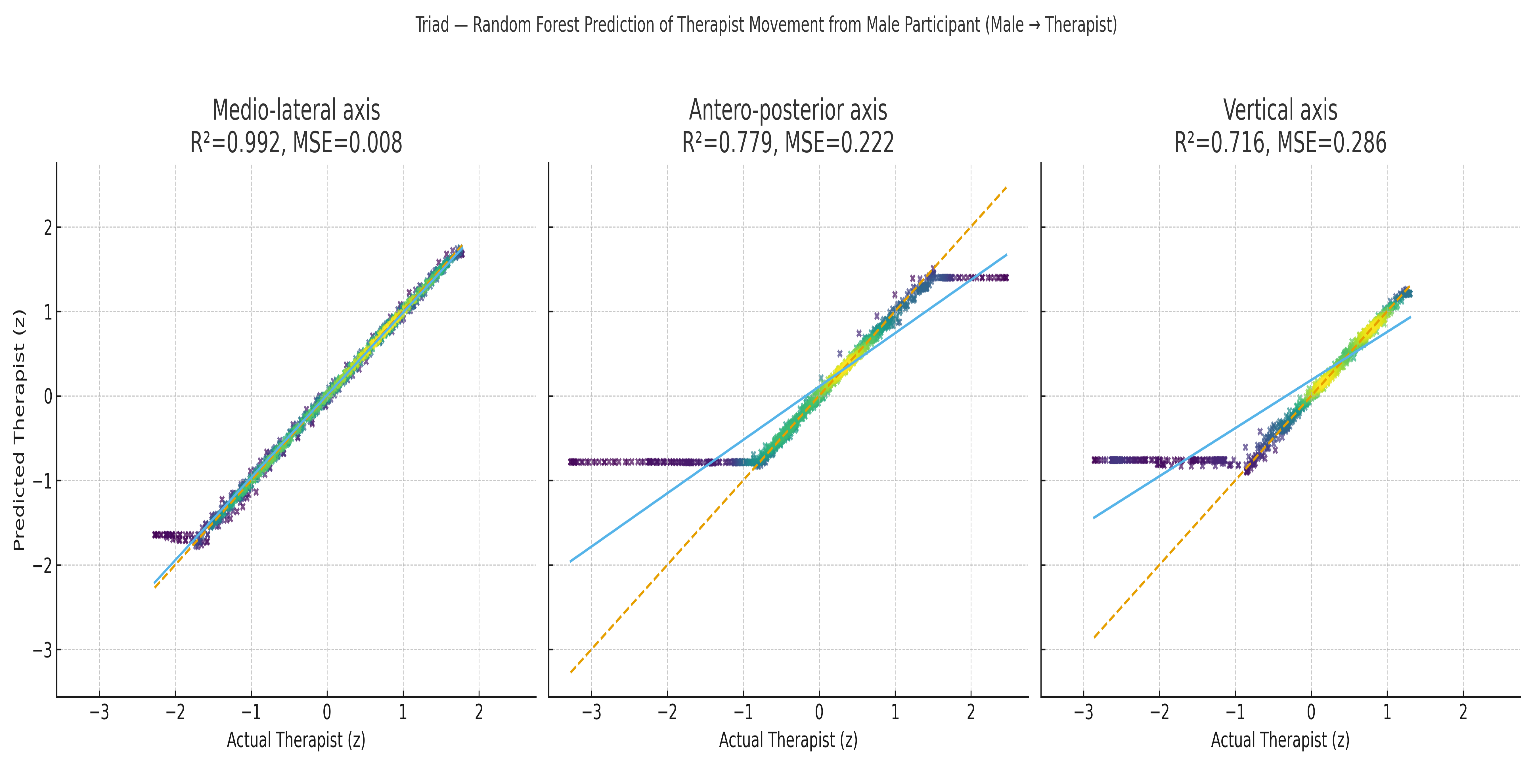

***Figure S16.*** *Random Forest regression: predicted vs. observed therapist movements (triadic condition; male client). Scatterplots show out-of-sample predictions of therapist centre-of-mass trajectories from the client’s lagged dynamics (eight lags; three-block time-ordered cross-validation). Points clustering along the identity line indicate high predictive accuracy in the medio–lateral axis and moderate accuracy in the antero–posterior and vertical axes, consistent with the reported R² and MSE values.*

**S4.10 Triadic Mirroring with Male Client — Granger Causality Analysis**

Granger causality analyses revealed no significant directional effects in the horizontal planes (see also Figure S17).

Along the ML axis, neither direction was significant (male → therapist: F(6, 991) = 0.96, p = .449; therapist → male: F(7, 988) = 0.55, p = .796).

Similarly, no significant effects were observed in the AP axis (male → therapist: F(7, 988) = 1.49, p = .168; therapist → male: F(8, 985) = 1.25, p = .267).

In the V axis, bidirectional Granger effects were detected (male → therapist: F(7, 988) = 2.42, p = .019; therapist → male: F(7, 988) = 2.34, p = .023).

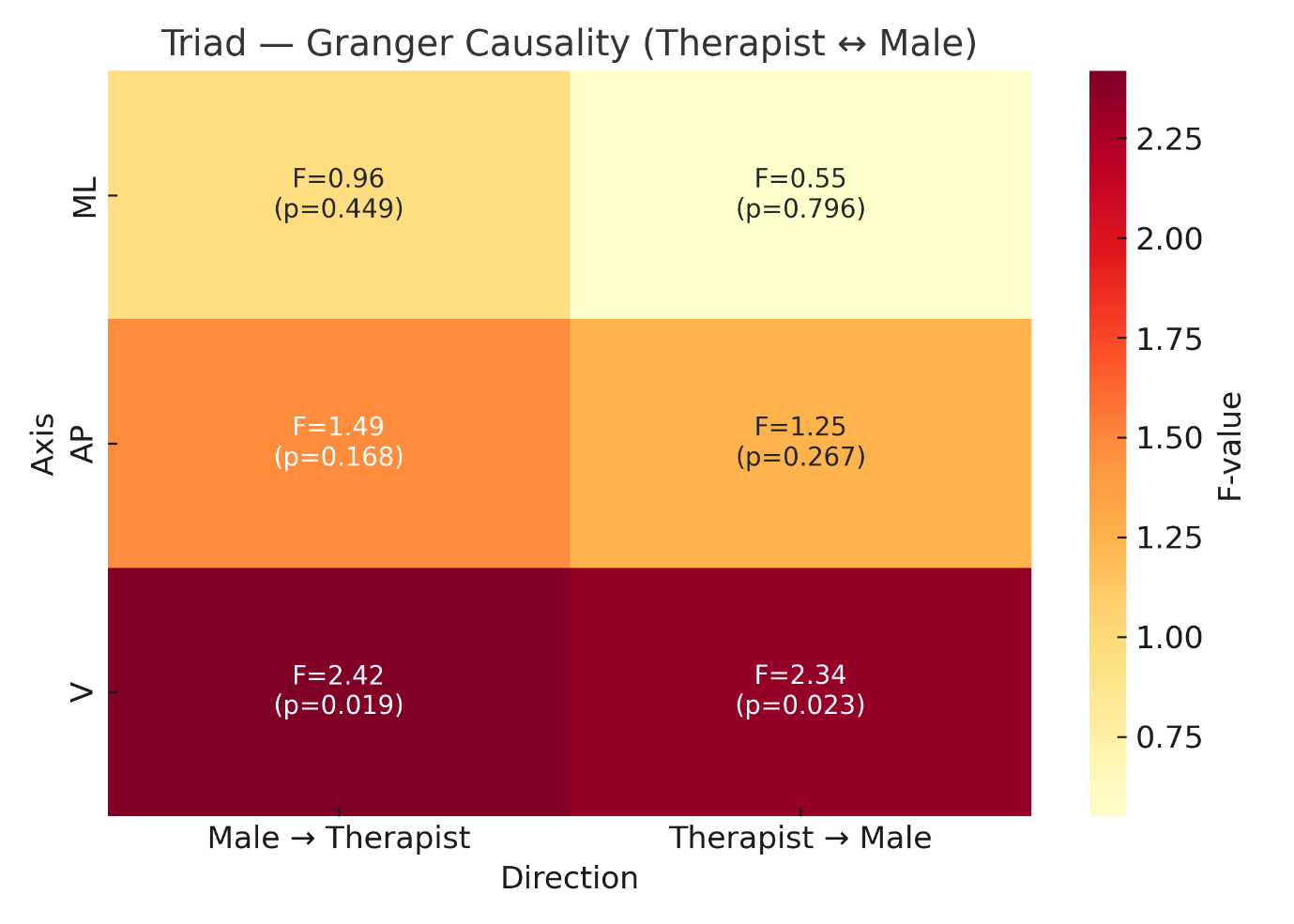

***Figure S17.*** *Granger causality heatmap for therapist and male client in the triadic task. Cells display F-values and p-values for each direction across ML, AP, and V axes, revealing significant bidirectional influence only in the vertical axis.*

**S4.11 Multivariate Modelling and Recurrence Analysis**

To extend beyond pairwise analyses and capture the therapist’s selective attunement to each client within the triadic interaction, we estimated axis-specific multivariate Vector Autoregressive (VAR) models. This approach complements the cross-correlation, phase synchronisation, Random Forest, and Granger causality analyses by simultaneously modelling the temporal dependencies among the therapist, female client, and male client within a single multivariate framework. In the context of Dance Movement Therapy, this allows assessment of whether the therapist’s moment-to-moment movement dynamics were preferentially driven by one client when attending to two individuals concurrently.

Separate VAR models were estimated for the medio–lateral (ML), antero–posterior (AP), and vertical (V) centre-of-mass (CoM) trajectories. For each axis, z-standardised CoM time series for the therapist, female client, and male client were entered simultaneously as endogenous variables. Model order selection based on information criteria consistently supported a VAR (7) specification across axes (see also Figure S18).

Across all three axes, the therapist’s movement exhibited strong autoregressive structure, indicating substantial self-dependence across successive lags. Focusing on the vertical axis—where earlier analyses revealed the strongest dyadic and triadic coupling—the female client’s prior movement did not significantly predict the therapist’s subsequent motion at lag 1 (β = −0.034, SE = 0.061, t = −0.55, p = .585) or at higher lags. Similarly, the male client’s prior movement did not exert a significant predictive influence on the therapist’s vertical dynamics (lag 1: β = −0.014, SE = 0.050, t = −0.29, p = .774). Across all seven lags, neither client’s coefficients reached statistical significance, whereas therapist self-dependence remained dominant.

Comparable patterns were observed in the ML and AP axes, where therapist movement was likewise primarily predicted by its own past dynamics, with no significant cross-client effects. Model diagnostics confirmed adequate fit and stability across axes: Ljung–Box tests indicated no residual autocorrelation at lag 10 (Q(10) = 6.37, p = .784), residuals were normally distributed (Jarque–Bera = 1.73, p = .421), and all eigenvalues of the companion matrix lay within the unit circle.

Taken together, these results indicate that, unlike in dyadic interactions, therapist movement in the triadic context was predominantly dominated by autoregressive structure across all movement dimensions, with no evidence of systematic temporal entrainment to either client. This suggests that when attending to two clients simultaneously, the therapist maintained a more autonomous movement trajectory, potentially reflecting the increased cognitive–motor demands of managing dual interpersonal engagements.

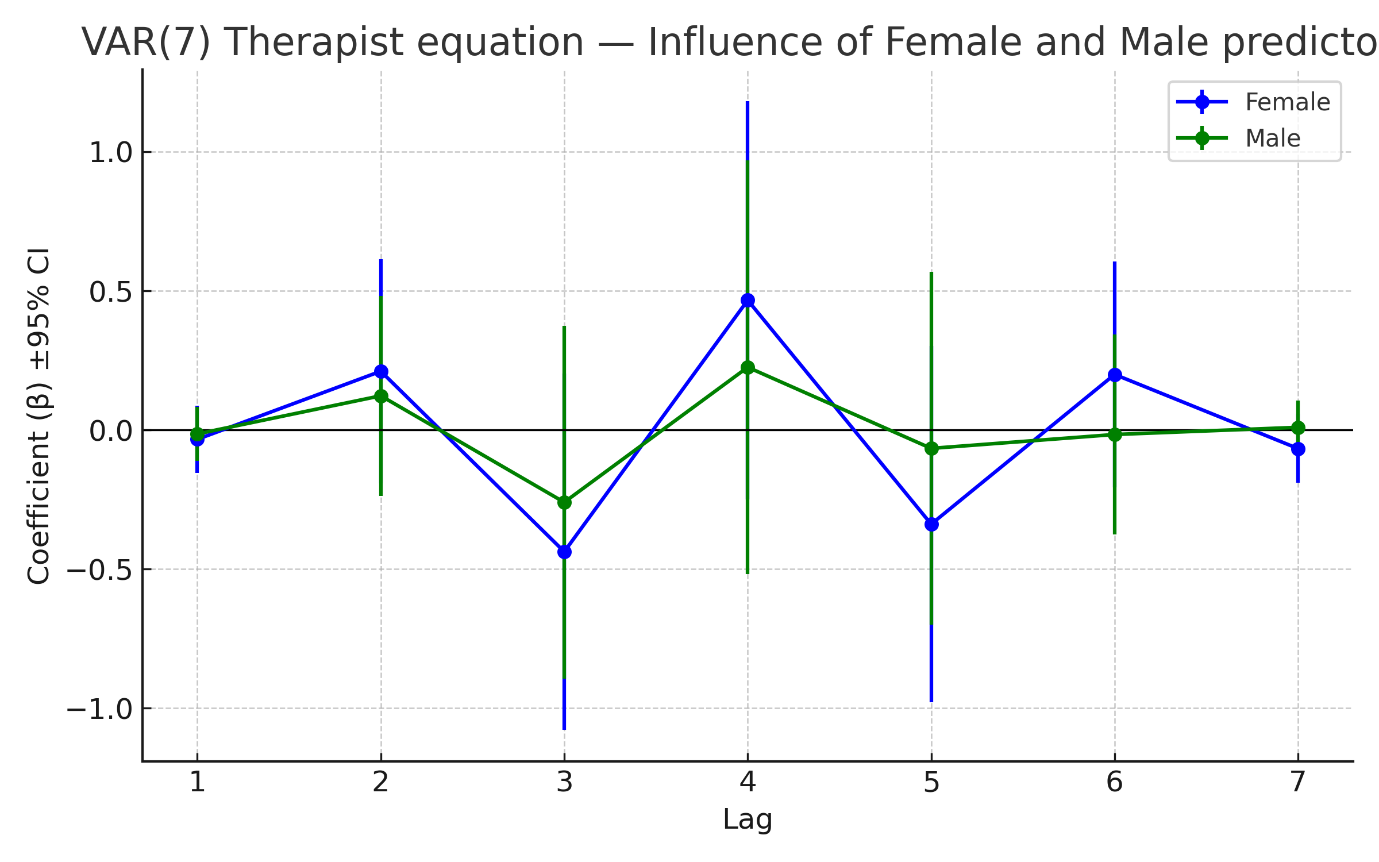

***Figure S18.*** *Coefficients (β ± 95% CI) from the VAR(7) model of therapist movement in the vertical axis, showing the estimated influence of female (blue) and male (green) clients across lags. None of the coefficients differ significantly from zero, indicating no reliable predictive effect from either client on the therapist’s movement.*

To complement the temporal dependency modelling afforded by VAR, joint recurrence quantification analysis (jRQA) was conducted to characterise the structural organisation of shared movement states between the therapist and each client in the triadic condition. Consistent with prior analyses indicating axis-specific effects, jRQA was applied to z-standardised vertical centre-of-mass (CoM) trajectories, as the vertical dimension showed the most consistent evidence of coupling across dyadic and triadic analyses (see also Figures S19 & S20).

Joint recurrence matrices were constructed using an absolute distance threshold corresponding to the 10th percentile of pairwise distances, and recurrence plots were generated separately for the therapist–male and therapist–female dyads. From these plots, standard recurrence metrics were extracted, including joint recurrence rate (JRR), determinism (DET), and laminarity (LAM), to quantify the density and temporal structure of shared movement states.

The therapist–male dyad exhibited a joint recurrence rate (JRR) of 0.0143, indicating that approximately 1.4% of time points corresponded to shared kinematic states. The determinism index (DET = 0.882) indicated that the majority of recurrence points formed diagonal line structures, reflecting temporally organised shared dynamics. The laminarity index (LAM = 0.949) further suggested that a large proportion of recurrence points were organised into vertical line structures, consistent with extended dwell times in similar movement states.

The therapist–female dyad showed a JRR of 0.0137, DET of 0.864, and LAM of 0.937. While these values were broadly comparable to those observed in the therapist–male dyad, they were marginally lower, indicating slightly reduced recurrence density and temporal structuring of shared states. Given the descriptive nature of the present analysis and the absence of inferential testing, these differences are interpreted cautiously as subtle variations in the organisation of shared movement dynamics rather than as evidence of categorical differences between dyads.

Visual inspection of the recurrence plots (Figures S19 & S20) was consistent with these quantitative patterns. Both dyads exhibited diagonal and vertical line structures indicative of temporally structured recurrence, with the therapist–male dyad displaying slightly denser and more sustained patterns than the therapist–female dyad, in line with the observed differences in DET and LAM.

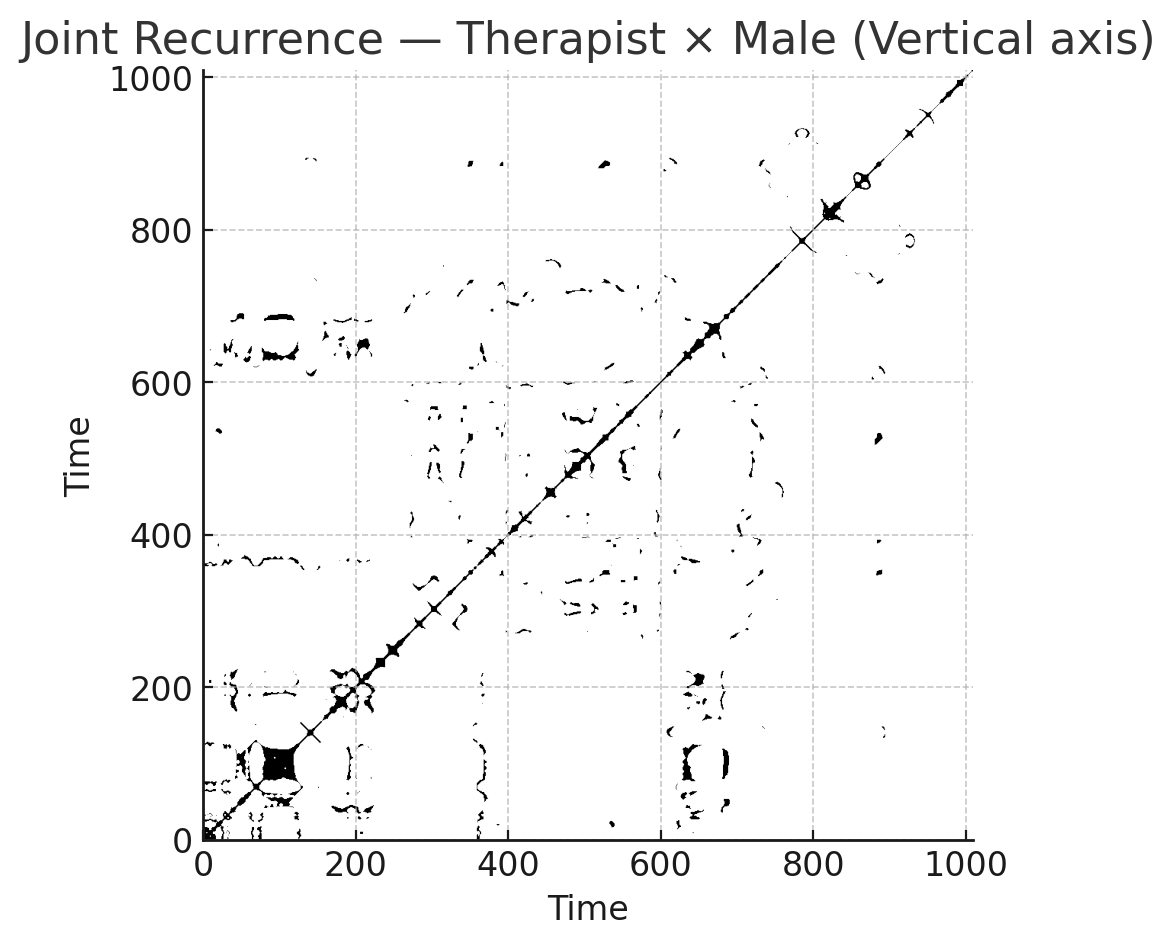

***Figure S19.*** *Recurrence plot of joint states between therapist and male client in the triadic task (vertical axis). Diagonal structures indicate deterministic coupling, while vertical patterns reflect laminar synchrony.*

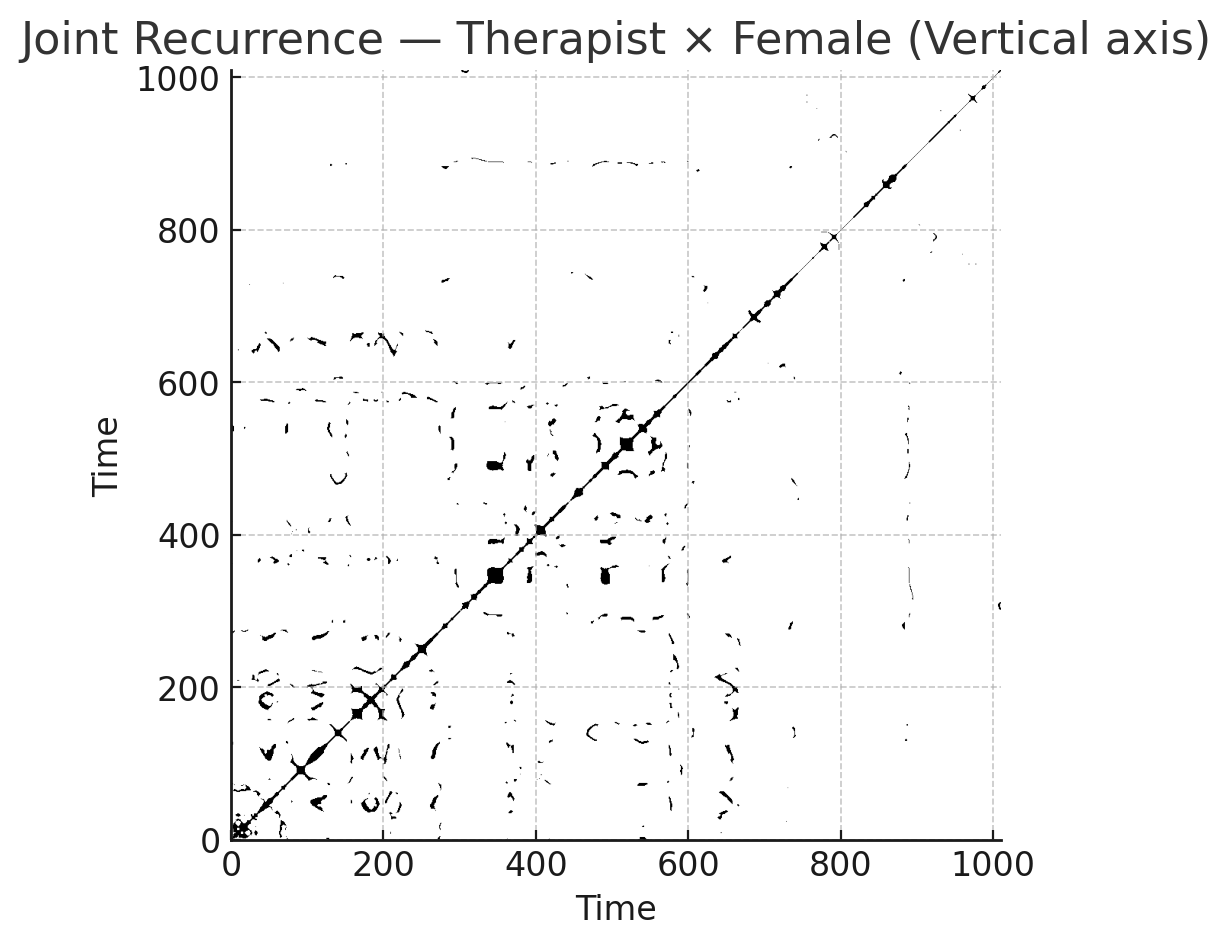

***Figure S20.*** *Recurrence plot of joint states between therapist and female client in the triadic task (vertical axis). Recurrence density and structure are marginally lower than in the therapist–male dyad.*
